# Allelic and Gene Dosage Effects Involving Uromodulin Aggregates Drive Autosomal Dominant Tubulointerstitial Kidney Disease

**DOI:** 10.1101/2022.09.13.507770

**Authors:** Guglielmo Schiano, Jennifer Lake, Marta Mariniello, Céline Schaeffer, Marianne Harvent, Luca Rampoldi, Eric Olinger, Olivier Devuyst

**Affiliations:** Mechanisms of Inherited Kidney Disorders, Institute of Physiology, University of Zurich; Zurich, CH-8057, Switzerland; Molecular Genetics of Renal Disorders, Division of Genetics and Cell Biology, IRCCS San Raffaele Scientific Institute, Milan, 20132, Italy; Institut de Recherche Expérimentale et Clinique, UCLouvain, Brussels, Belgium; Translational and Clinical Research Institute, Newcastle University; Newcastle upon Tyne, NE1 3BZ, United Kingdom; Center for Human Genetics, Cliniques Universitaires Saint-Luc, UCLouvain, Brussels, Belgium

## Abstract

Missense mutations in the *UMOD* gene encoding uromodulin cause autosomal dominant tubulointerstitial kidney disease (ADTKD), one of the most common monogenic kidney diseases. A pressing need for ADTKD is to bridge the gap between postulated gain-of-function mutations and organ damage - a prerequisite for therapeutic development. Based on two missense *UMOD* mutations associated with divergent progression of ADTKD, we generated *Umod*^*C171Y*^ and *Umod*^*R186S*^ knock-in mice that showed strong allelic and gene dosage effects, with distinct dynamic pathways impacting on uromodulin trafficking, formation of intracellular aggregates, activation of ER stress, unfolded protein and immune responses, kidney damage and progression to kidney failure. Deletion of the wild-type *Umod* allele in heterozygous *Umod*^*R186S*^ mice increased the formation of uromodulin aggregates and ER stress, indicating a protective role of wild-type uromodulin. Studies in kidney tubular cells confirmed biochemical differences between distinct uromodulin aggregates, with activation of specific quality control and clearance mechanisms. Enhancement of autophagy by starvation and mTORC1 inhibition decreased the uromodulin aggregates, suggesting a therapeutic strategy. These studies substantiate a model for allelic effects and the role of toxic aggregates in the progression of ADTKD-*UMOD*, with relevance for toxic gain-of-function mechanisms and for strategies to improve clearance of mutant uromodulin.

## INTRODUCTION

Autosomal dominant tubulointerstitial kidney disease (ADTKD) includes a set of rare kidney disorders characterized by tubular damage and interstitial fibrosis in the absence of glomerular lesions. Affected individuals develop chronic kidney disease (CKD) that ultimately progresses to kidney failure. There is currently no specific treatment to alter the progression of ADTKD (1). Mutations in several genes can cause ADTKD, the most common affecting the *UMOD* gene (ADTKD-*UMOD*), which is exclusively expressed in the kidney and encodes uromodulin, the most abundant protein excreted in normal urine (2). ADTKD-*UMOD* (MIM #162000) is one of the most common monogenic kidney diseases, with an overall prevalence of ∼2% in patients with kidney failure (3, 4).

Uromodulin is a glycosylphosphatidylinositol (GPI)-anchored glycoprotein, produced mainly by the cells lining the thick ascending limb (TAL) of the loop of Henle (5). After the formation of 24 disulfide bridges and extensive glycosylation, uromodulin is trafficked to the apical plasma membrane, where it is cleaved and released into the tubular lumen to form high-molecular-weight polymers (6). Uromodulin exerts important roles in tubular cells and lumen, including regulation of sodium transport (7, 8), inhibition of kidney stone formation (9) and protection against urinary tract infections (10, 11).

More than 125 ADTKD-*UMOD* mutations have been reported, essentially missense mutations that replace or introduce a cysteine residue (1, 2). Analyses of kidney biopsies from patients with ADTKD-*UMOD* revealed a striking accumulation of uromodulin in the endoplasmic reticulum (ER) of TAL cells, suggestive of ER storage disease (12-14). Consistent with abnormal processing, a dramatic reduction in uromodulin levels in the urine was observed in individuals with ADTKD-*UMOD* (12, 15). In vitro analyses confirmed that mutations in uromodulin cause defective ER to Golgi transport, with activation of the unfolded protein response (UPR) and a suggestive correlation between the severity of the trafficking defect of uromodulin mutants and the age of ESKD (16, 17).

Mouse models carrying *Umod* mutations have been generated using ENU mutagenesis, transgenesis, homologous recombination or CRISPR/Cas9 (18-22). These mice recapitulate the key features of ADTKD-*UMOD*, including uromodulin accumulation and tubulointerstitial damage, activation of ER stress pathways and progressive CKD. However, the limited information coming from these models and the unknown consequences of various types of mutant uromodulin accumulation leave open the mechanisms of disease progression — preventing therapeutic developments. These issues are emphasized by the strong interfamilial variability in the progression of ADTKD-*UMOD* (2, 23) and a possible gene-dosage effect on uromodulin processing (24). Wild-type uromodulin may also play a role, analogous to other dominant, gain-of-toxic function disorders such as Huntington disease, where the lack of wild-type huntingtin causes a more severe phenotype (25, 26).

To elucidate the gap between postulated gain-of-function mutations and clinically relevant endpoints, we selected two missense *UMOD* mutations representative of divergent disease progression in patients with ADTKD-*UMOD*. We generated *Umod* knock-in (KI) mice carrying these p. C170Y and p. R185S *UMOD* mutations and investigated allelic and gene dosage effects on the formation of intracellular aggregates, the role of wild-type uromodulin and the downstream signaling pathways leading to CKD. Analyses in stably transduced kidney tubular cells revealed distinct ER accumulation of premature mutant uromodulin and activation of the ER stress response, with the formation of biochemically distinct aggregates and differential activation of clearing mechanisms. These studies decipher the pathways linking the formation of intracellular uromodulin aggregates, unfolded protein response and kidney damage in ADTKD, with relevance for therapeutic strategies and for toxic gain-of-function mechanisms in autosomal dominant diseases.

## RESULTS

### Identification of *UMOD* mutations associated with divergent ADTKD progression

We characterized missense mutations associated with divergent severity of ADTKD-*UMOD* as a first step to dissect properties of uromodulin mutants and pathways driving kidney disease. A review of 12 missense *UMOD* mutations detected in at least 2 genetically confirmed ADTKD-*UMOD* individuals reaching kidney failure in the Belgo-Swiss ADTKD Registry (2), confirmed a strong influence of the mutation on age at kidney failure (test for linear trend: p<0.0001) (Figure 1A). Two clusters of mutations associated with earlier (before ∼40 years) and later (50-69 years) onset of kidney failure were identified. Two representative mutations, p. Arg185Ser (earlier-onset) and p. Cys170Tyr (later-onset), were selected since they: (i) were the most prevalent in both clusters; (ii) involved cysteine and noncysteine residues, potentially relevant for toxic functions; and (iii) were detected in typical ADTKD-*UMOD* families with available kidney biopsy. Patients with the *UMOD* p. Arg185Ser mutation showed a much faster disease progression than those with the p. Cys170Tyr mutation (N=9 in each group; median age of kidney failure: 42 years vs. 69 years), clearly distinct from the aggregated data for 60 ADTKD patients harboring 22 other *UMOD* mutations in the Registry (median age of kidney failure: 56 years; log rank test: p=0.0035; Figure 1B). Both mutations segregated in typical multiplex ADTKD families and were found to be (likely) pathogenic by in silico analyses (Figure 1C; <u>Supplementary Tables 1-3</u>). Examination of kidney biopsies revealed that the p. Arg185Ser mutation was associated with mutant uromodulin ER accumulation, ER stress and widespread fibrosis, whereas the p. Cys170Tyr uromodulin was predominantly detected on the apical plasma membrane, without induction of ER stress and no detectable fibrosis (Figure 1D).

**Figure 1.**
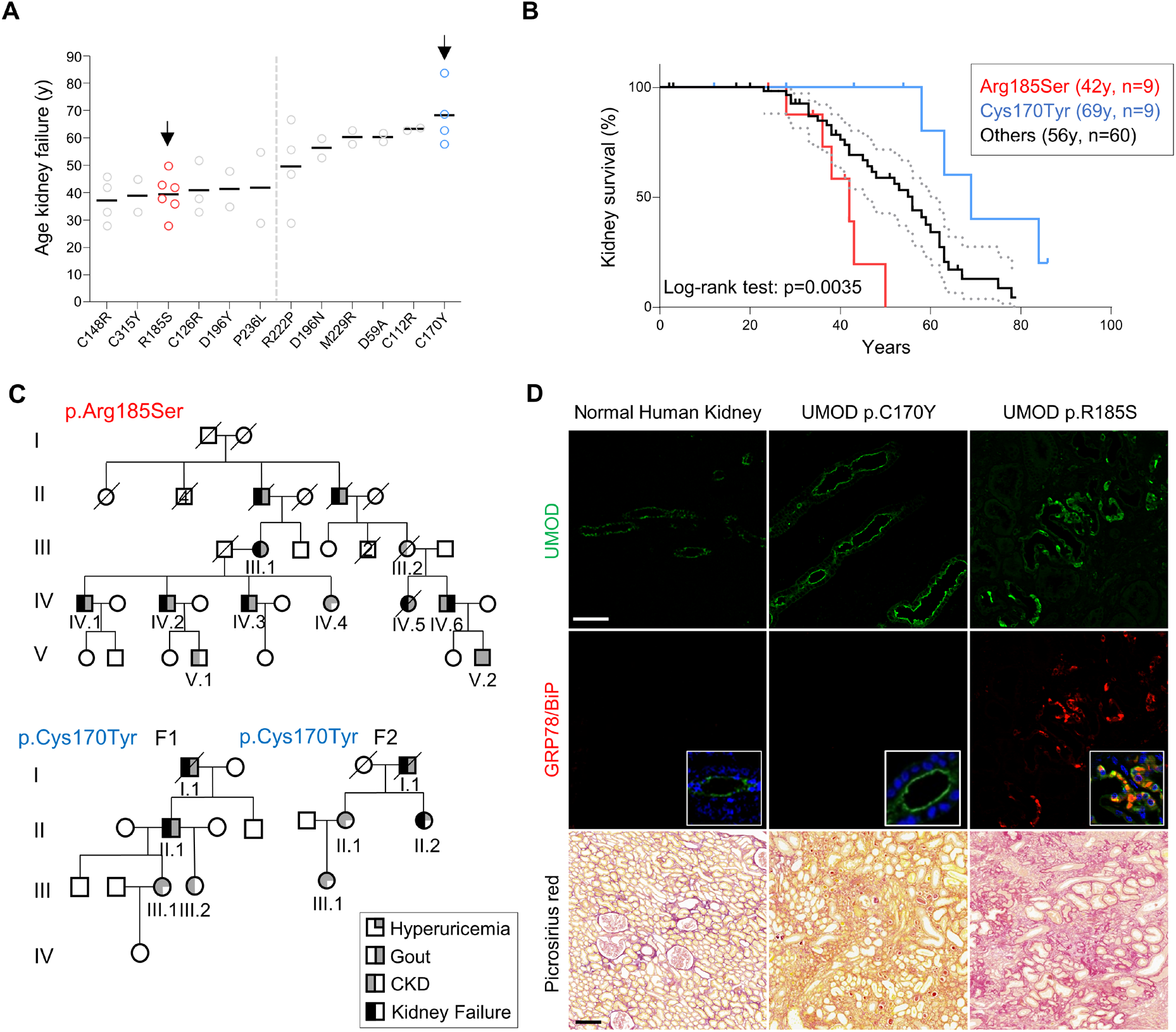
Mutations associated with divergent progression of ADTKD-*UMOD*. (**A**) Age at onset of kidney failure for ADTKD-*UMOD* patients with the indicated *UMOD* mutations. Only mutations with at least 2 individuals reaching kidney failure at documented age are represented here. Two clusters of *UMOD* mutations associated with earlier-onset and later-onset kidney failure with two cluster-representative mutations (arrows, see result text) are highlighted. (**B**) Kaplan-Meier curve of kidney survival in patients with the *UMOD* p. Arg185Ser mutation (*N*=9, median age at kidney failure: 42 years), the *UMOD* p. Cys170Tyr mutation (*N*=9, median age at kidney failure: 69 years) and 60 ADTKD-*UMOD* patients from the Belgo-Swiss registry (see Material & Methods) with 24 different *UMOD* mutations (median age at kidney failure: 56 years). A log-rank test was used for comparison of survival curves. (**C**) Pedigrees of three multiplex families with ADTKD in which representative *UMOD* mutations p. Arg185Ser and p. Cys170Tyr have been identified. Females are represented by circles and males by squares, and phenotypes are denoted as indicated. Clinical features are detailed for each patient in Supplementary Tables 1-2. (**D**) Representative confocal analysis of uromodulin (UMOD, green), GRP78/BiP (red) and Picrosirius Red staining of kidney nephrectomy samples from ADTKD-*UMOD* patients (p. Cys170Tyr – F1, II.1; p. Arg185Ser - IV.5). For immunofluorescence, nuclei were counterstained with DAPI (blue). Scale bar: 25 μm (top), 100 μm (bottom).

Thus, based on kidney failure, we identified two *UMOD* missense mutations representative of rapid- and slow-progressing ADTKD-*UMOD* with potentially divergent pathomechanisms, as suggested from kidney tissue analysis.

### *Umod* KI mice show strong allelic and gene dosage effects on uromodulin processing and kidney damage

Based on the representative *UMOD* mutations (C170Y and R185S), we generated two KI mouse models harboring the equivalent *Umod* mutations (C171Y and R186S; Figures 2A, B). These *Umod* KI mice were born in the expected Mendelian ratio and were viable and fertile. We first verified whether the *Umod* mutations were reflected by maturation and trafficking defects (Figures 2C, D). The mature, fully glycosylated uromodulin band (∼100 kDa) was detected with variable intensity in *Umod*^C171Y^ kidneys (Figure 2C). In homozygous *Umod*^C171Y/C171Y^ kidneys, the uromodulin signal included the mature form and additional bands at ∼80 kDa, potentially corresponding to immature isoforms carrying ER-type glycans, and high-molecular-weight (HMW, ∼160-200 kDa) bands, compatible with uromodulin aggregates (Figure 2C). In contrast, the ∼100 kDa band corresponding to mature uromodulin was not detected in the *Umod*^R186S^ kidneys, whereas strong signals for immature (∼80 kDa) and HMW aggregate forms of uromodulin, were detected at 1 month of age (Figure 2D). Treatment with endoglycosidase H (Endo H) or peptide:N-glycosidase F (PNGase F) confirmed the presence of immature forms of uromodulin carrying ER-type glycosylation in both homozygous *Umod*^C171Y^ and *Umod*^R186S^ kidney samples (<u>Supplementary Figures 1A, B</u>). The changes in uromodulin maturation impacted its urinary excretion: striking reductions were observed in *Umod*^R186S^ mice (Figure 2D), whereas urinary levels were unchanged in *Umod*^C171Y/+^ mice, being only decreased in the *Umod*^C171Y/C171Y^ mice (Figure 2C).

**Figure 2.**
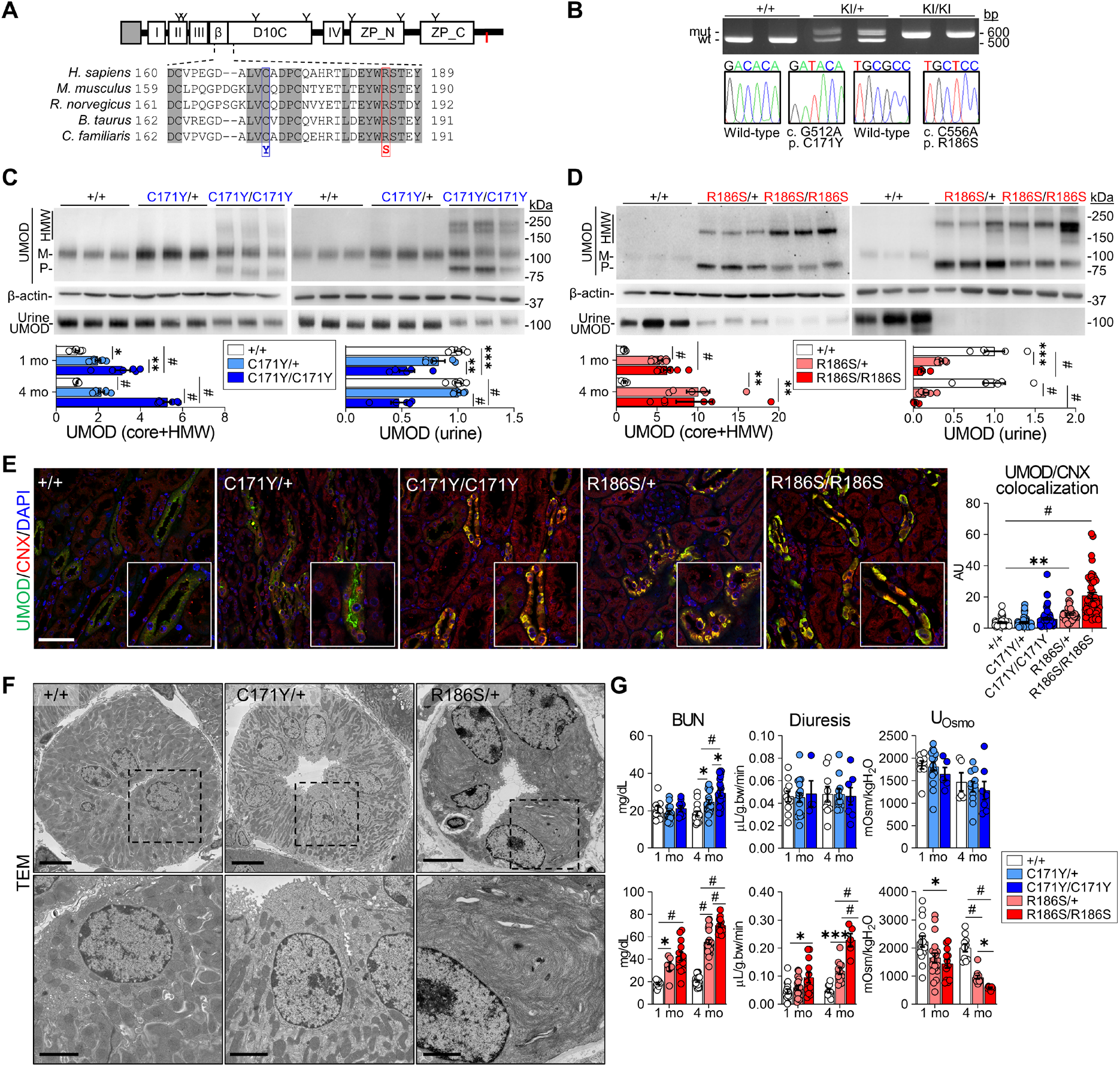
Characterization of the *Umod*^C171Y^ and *Umod*^R186S^ mouse models. (**A**) Uromodulin (UMOD) domain architecture including a signal peptide (gray box), four EGF-like domains (I, II, III, IV), a beta-hairpin (β), a cysteine-rich domain (D10C) and a bipartite zona pellucida domain (ZP_N, ZP_C). The GPI-anchoring site (red line) and glycosylation sites (Y) are indicated. Alignment of uromodulin amino acid sequences in different mammalian species is shown, with identity regions displayed as shadowed. Mutations of interest are indicated in red (p. R186S) and blue (p. C171Y). (**B**) Genotyping and sequencing chromatogram of *Umod* KI mice, showing the regions surrounding the two mutations (c. C512, c. C556) in *Umod*. (**C, D**) Immunoblot analysis of kidney and urine uromodulin (UMOD) in 1- and 4-month-old *Umod*^C171Y^ (**c**) and *Umod*^R186S^ (**D**) mice. For kidney lysates, β-actin was used as a loading control. Urine samples were normalized to creatinine concentration (*n*=5 to 9 animals per group). Densitometry analysis is relative to *Umod*^+/+^. M: mature; P: precursor; HMW: high molecular weight. The mature and precursor uromodulin are collectively referred to as the core. (**E**) Representative immunofluorescence analysis of uromodulin (UMOD, green) and calnexin (CNX, red) in kidney sections from 4-month-old mice (*n*=47 to 70 tubules per group). Nuclei are counterstained with DAPI (blue). Scale bar: 25 μm. (**F**) Transmission electron microscopy (TEM) of kidney sections from 3-month-old *Umod* mice, showing progressive ER expansion and hyperplasia. Squares indicate images shown at higher magnification. Scale bar: 5 μm for low magnification, 2 μm for high magnification. (**G**) Main clinical parameters of *Umod*^C171Y^ and *Umod*^R186S^ mice at 1 and 4 months (*n*=5 to 16 animals per group). Bars indicate the mean ± SEM. One-way ANOVA with Tukey’s post hoc test; ^*^*P <* 0.05, ^**^*P* < 0.01, ^***^*P <* 0.001, ^#^*P <* 0.0001.

We next analyzed whether specific mutations affect the distribution of uromodulin in the *Umod* KI kidneys. Dual labeling with the ER marker calnexin showed a mild ER retention of uromodulin with residual apical signal in *Umod*^C171Y^ kidneys, contrasting with a major ER retention of uromodulin and lack of apical signal in the *Umod*^R186S^ kidneys (Figure 2e), mimicking the human situation (Figure 1D). At the subcellular level, the massive accumulation of uromodulin caused expansion and hyperplasia of the ER in *Umod*^R186S/+^ kidneys, as shown by electron microscopy (Figure 2F) and correlative light-electron microscopy (CLEM) analysis (<u>Supplementary Figure 1C</u>). The intracellular uromodulin aggregates in *Umod*^R186S/+^ kidneys were already observed at 2 weeks of age (<u>Supplementary Figures 2A-C</u>), with only faint signals for mature uromodulin (<u>Supplementary Figure 2B</u>) and apical staining in TAL cells (<u>Supplementary Figure 2C)</u> in these samples.

To test whether the changes in uromodulin processing affected disease progression, we analyzed the plasma and urine biochemistry of *Umod* KI mice compared to wild-type control littermates. The *Umod*^C171Y^ mice showed a progressive increase in BUN at 4 months, whereas the *Umod*^R186S^ mice showed significantly increased BUN levels at 1 month, with a strong urinary concentration defect, reduced fractional excretion of uric acid, and inappropriate urinary calcium excretion (Figure 2G; <u>Supplementary Tables 4, 5</u>).

Collectively, these data indicate that the two *Umod* KI lines present typical manifestations of ADTKD-*UMOD*, with strong allelic and gene-dosage effects on uromodulin processing and severity of kidney damage, in line with observations in humans.

### Distinct mutational effects on the properties of uromodulin aggregates

To test the impact of the biochemical nature of uromodulin aggregates on the allelic effects, we first analyzed kidney uromodulin in its native form and subsequently performed partial and total protein denaturation (Figure 3A). Under native conditions, *Umod*^C171Y^ lysates showed an additional higher band, which was not observed in *Umod*^+/+^ kidneys. Moreover, part of the uromodulin from *Umod*^C171Y/C171Y^ and most in *Umod*^R186S^ lysates migrated even less efficiently (Figure 3B). Following partial denaturation, *Umod*^C171Y/+^ kidney lysates showed only the band corresponding to mature uromodulin, suggesting that aggregates in these samples were mediated by weak interactions. In contrast, HMW bands were still detected in *Umod*^C171Y/C171Y^ and *Umod*^R186S^ kidneys, which could only be resolved when using reducing conditions, suggesting the involvement of intermolecular disulfide bridges (Figure 3B). The solubility of uromodulin aggregates was assessed using a soluble/insoluble fractionation protocol (Figure 3C): wild-type uromodulin was exclusively localized in the soluble fraction, whereas premature forms and HMW uromodulin aggregates were mainly in insoluble fractions (Figure 3D). Lysates from both homozygous mutant kidneys showed an enriched signal in the insoluble fraction compared to heterozygotes. Although no signal for premature or HMW uromodulin was detected in *Umod*^C171Y/+^ kidneys, mature uromodulin was detected in both soluble and insoluble fractions. This suggests that p.C171Y uromodulin is characterized by a partially tolerated folding defect and is still undergoing complete maturation. However, once the mutant protein reaches the apical plasma membrane, misfolding may reoccur, causing increased retention.

**Figure 3.**
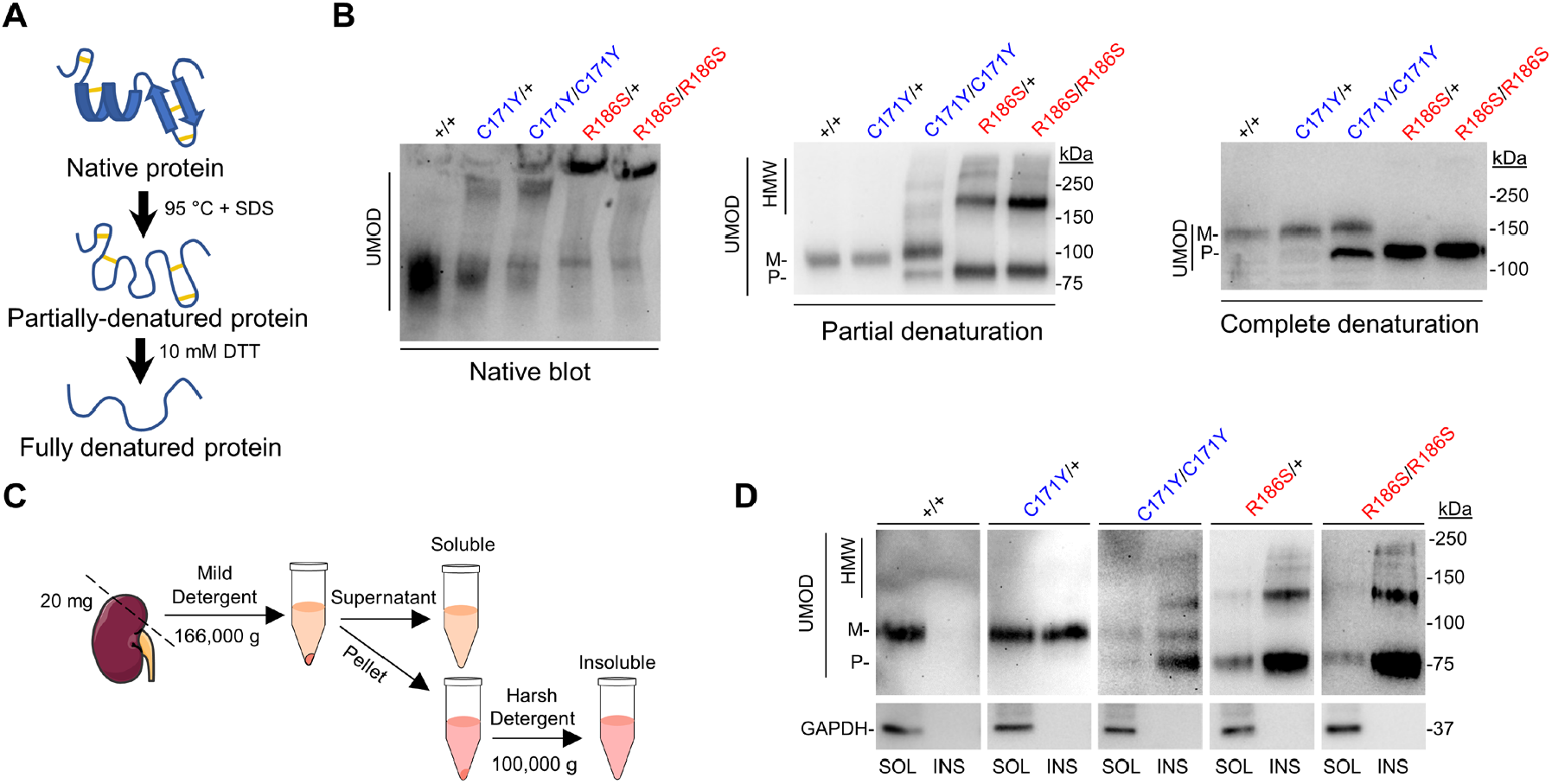
Biochemical profiling of high molecular weight uromodulin aggregates. (**A**) Schematic representation of the partial and total protein denaturation treatments. (**B**) Immunoblot analysis of kidney lysates from *Umod* KI mice in native (left), partially denatured (center) and completely denatured (right) conditions, showing the sample treatment-dependent effect on uromodulin migration. (**C**) Schematic representation of the solubility assay protocol. (**D**) Solubility assay of uromodulin in kidney lysates from 4-month-old *Umod* KI mice. GAPDH was used as a purity marker for the insoluble fraction.

These data show that both weak and strong interactions may participate in the formation of aberrant uromodulin aggregates, whose biochemical properties strongly depend on the mutation involved.

### Wild-type uromodulin protects against mutant uromodulin aggregation

Given the gene-dosage effect observed in both *Umod* KI models and the early-onset disease in *Umod*^R186S^ mice, we wondered whether the increased severity observed in *Umod*^R186S/R186S^ mice was solely due to an increased load of mutant uromodulin, or whether the wild-type allele may attenuate the phenotype. To this end, we crossed *Umod*^R186S/+^ with *Umod*^*-*/*-*^ mice to obtain *Umod*^R186S/*-*^ mice, with the expected decrease in *Umod* mRNA (Figures 4A, B) and in the urinary levels of uromodulin (Figure 4C). Despite the decreased global expression of uromodulin and comparable protein levels between *Umod*^R186S/*-*^ and *Umod*^R186S/+^, the former showed an increase in HMW aggregates, and a pronounced reduction in premature uromodulin, was observed in *Umod*^R186S/*-*^ kidneys compared to *Umod*^R186S/+^ kidneys (Figure 4D). These differences were confirmed in isolated TAL segments from *Umod*^R186S^ kidneys, where deletion of the wild-type allele increased the expression of the ER-resident chaperone GRP78 (Figure 4E). The impact of HMW uromodulin accumulation in the *Umod*^R186S/*-*^ kidneys was further evidenced by a strong upregulation in inflammation (*Adgre1, Cd68, Ptrpc, Tlr4*), fibrosis (*Fn1, Tgfb*1) and tubular damage (*Lcn2*) markers compared to *Umod*^+/+^ and *Umod*^R186S/+^ kidneys (Figure 4F), and by larger numbers of CD3^+^ (lymphocyte) cells close to uromodulin-positive TAL segments (Figure 4G). Of note, these changes in *Umod*^R186S/*-*^ compared to *Umod*^R186S/+^ mice were not reflected in terms of ER retention of uromodulin (Figure 4G) or by clinical and biological parameters (<u>Supplementary Table 6</u>).

**Figure 4.**
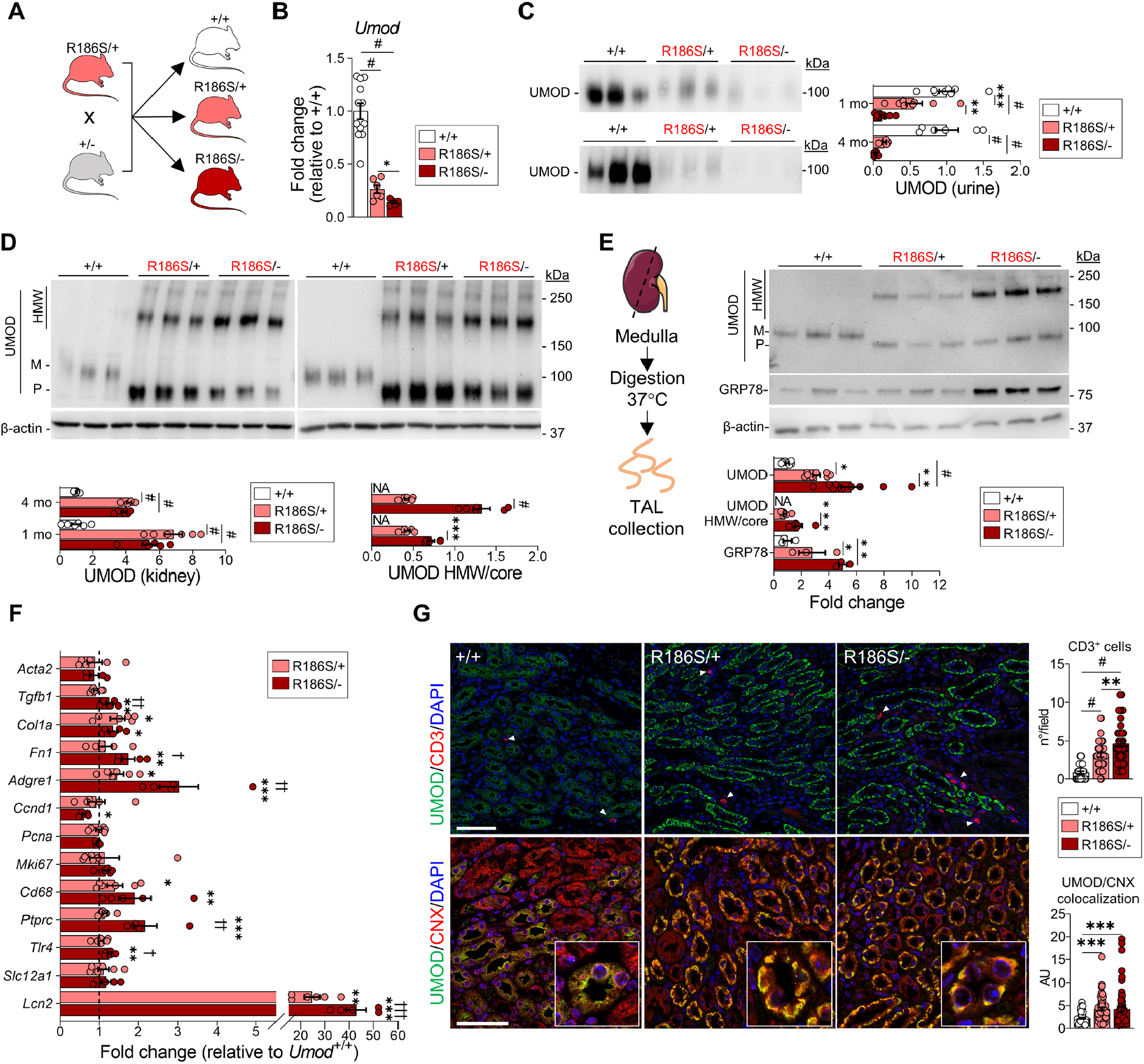
Deletion of wild-type uromodulin increases mutant uromodulin aggregation. (**A**) Strategy for the generation of *Umod*^R186S/-^ mice. (**B**) *Umod* transcript levels evaluated by RT-qPCR on total kidney extracts from 1-month *Umod*^R186S/+,^ *Umod*^R186S/*-*^, and *Umod*^+/+^ mice (*n*=5 to 12 animals per group). (**C**) Immunoblot analysis of urine uromodulin (UMOD) in 1- and 4-month *Umod*^R186S/*-*^ mice. Urine was loaded according to creatinine concentration (*n=*5 to 6 animals per group). (**D, E**) Representative immunoblot analysis of UMOD in whole kidney at 1 month or 4 months (*n=*5 to 6 animals per group) (**D**) or of UMOD and GRP78 in isolated TAL (*n*=3 to 9 TAL fractions per group). (**E**). β-actin was used as a loading control. M: mature; P: precursor; HMW: high molecular weight. The mature and precursor uromodulin are collectively referred to as the core. (**F**) Transcript levels of inflammation, proliferation, fibrotic markers and *Slc12a1* as internal control assessed by RT-qPCR on total kidney extracts from mice at 1 month. (*n*= 5 to 9 animals per group). (**G**) Confocal analysis of kidney sections from 1-month mice stained with anti-uromodulin (green) and anti-CD3 or anti-calnexin (CNX, red). *n* = 40 fields from 4 kidneys per condition (up), *n* ≥ 57 tubules from 3 kidneys per condition (down). Nuclei are stained with DAPI (blue). Scale bar: 50 μm. Bars indicate the mean ± SEM. One-way ANOVA with Tukey’s post hoc test, ^*^*P* < 0.05, ^**^*P* < 0.01, ^***^*P* < 0.001, ^#^*P* < 0.0001 compared to *Umod*^+/+, †^*P* < 0.05; ^††^*P* < 0.01, ^†††^*P* < 0.001 *Umod*^R186S/+^ versus *Umod*^R186S/-^.

### Differential activation of the UPR and kidney damage pathways by mutant uromodulin

The striking differences in the amount of biochemically distinct uromodulin aggregates in the *Umod* mouse lines (Figure 5A) led us to investigate their differential effects on the ER and UPR stress pathways. The signal for the UPR gatekeeper protein GRP78/BiP was low in the (uromodulin-positive) TAL segments of *Umod*^+/+^ and *Umod*^C171Y/+^ mice, reflecting normal uromodulin processing, but strikingly increased in *Umod*^C171Y/C171Y^, *Umod*^R186S/+^, *Umod*^R186S/-^ and *Umod*^R186S/R186S^ kidneys, consistent with uromodulin accumulation in the ER (Figure 5B). Immunoblot analyses showed that uromodulin storage upregulated the PERK (PERK, ATF4) and IRE1α branches of the UPR in *Umod*^R186S^ (<u>Supplementary Figure 3A, C-D</u>) but not in *Umod*^C171Y^ kidneys (<u>Supplementary Figure 3B</u>). Importantly, activation of PERK did not trigger apoptosis, as indicated by the lack of TUNEL-positive cells or increased cleavage of caspase-3 in the *Umod*^R186S^ kidneys (<u>Supplementary Figure 4</u>). These results indicate that the ER accumulation of specific mutant uromodulin triggers the UPR based on activation of the PERK and IRE1α branches.

**Figure 5.**
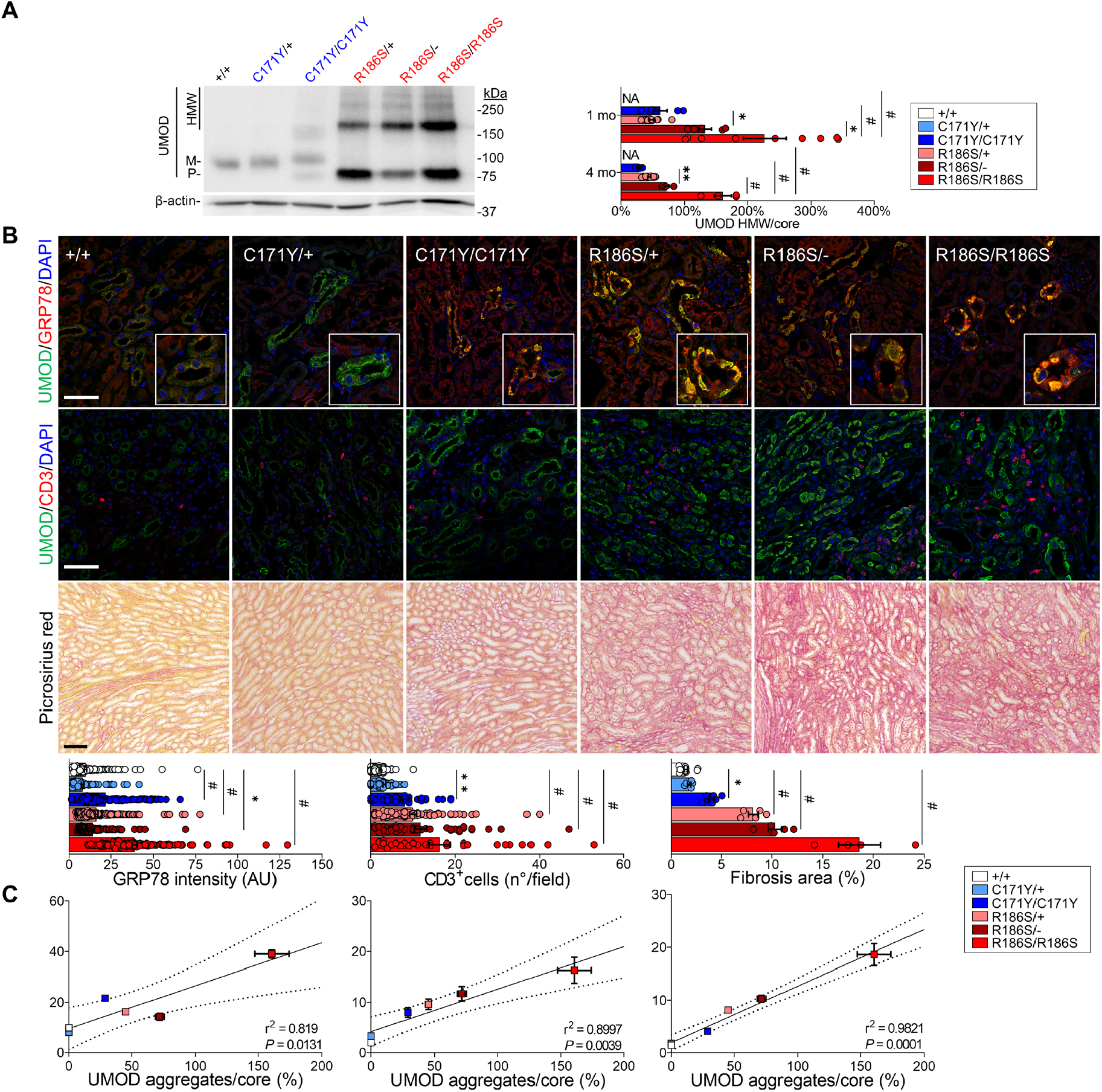
Mutant uromodulin aggregates drive severity of ADTKD-*UMOD* in mouse models. (**A**) Representative immunoblots for uromodulin in kidneys from 1-month wild-type and mutant mice, showing distinct patterns for mature, precursor and HMW bands. β-actin was used as a loading control (*n=*4 to 12 animals per group). M: mature; P: precursor; HMW: high molecular weight. The mature and precursor uromodulin are collectively referred to as the core. (**B**) Immunofluorescence analysis of uromodulin and GRP78 (top) or CD3 (middle) and picrosirius red staining on kidney sections from 4-month-old *Umod* KI mice (*n*=3 to 11 animals per group). Scale bar: 25 μm for IF, 50 μm for picrosirius red. Bars indicate the mean ± SEM. One-way ANOVA with Tukey’s post hoc analysis, ^*^*P* < 0.05, ^**^*P* < 0.01, ^#^*P* < 0.0001. (**C**) Linear regression (black line) illustrating the correlation between UMOD aggregates and GRP78 intensity (left), CD3^+^ infiltrates (middle) and interstitial fibrosis (right) in 4-month-old *Umod* KI mice. The dotted lines show the 95% confidence intervals. Dots indicate the mean ± SEM. The equations for the curves are y= 0.1695x + 9.567 (left), y = 0.08372x + 4.203 (middle), and y = 0.1073x + 1.915 (right).

Importantly, the different *UMOD* mutations induced distinct expression patterns for ER-phagy genes involved in protein quality control mechanisms. In particular, the expression levels of *Rtn3, Sec62* and *Ccpg1* were significantly higher in *Umod*^C171Y^ kidneys than in *Umod*^R186S^ kidneys, while *Retreg1* was downregulated in both *Umod*^R186S^ and *Umod*^C171Y/C171Y^ kidneys containing uromodulin aggregates (<u>Supplementary Figure 5A</u>). The activation of cellular degradation pathways (autophagy and proteasome) to mitigate misfolded protein toxicity was evidenced by the upregulation of SQSTM-1/p62 in *Umod* KI kidneys that paralleled the allelic and gene dosage effects on aggregate formation (<u>Supplementary Figure 5B</u>).

Tubulointerstitial damage and fibrosis characterize end-organ damage in ADTKD-*UMOD*. Analyses in age- and gender-matched *Umod* lines revealed that uromodulin accumulation was paralleled by a progressive infiltration of CD3+ cells and the onset of fibrosis, starting around the cortical TALs and later expanding to the outer medulla (Figure 5B). Quantitative analyses indicated tight correlations between the amount of uromodulin aggregates and the levels of GRP78 upregulation, the number of CD3+ cells and the extent of interstitial fibrosis in the kidneys of the different *Umod* lines, substantiating the role of uromodulin aggregates in driving interstitial inflammation and fibrosis (Figure 5C).

### Mutant uromodulin triggers distinct pathways impacting disease progression

Based on the strong allelic effects on the ER accumulation of uromodulin, formation of aggregates, and generation of ER stress, reflected by divergent disease progression in the R186S and C171Y *Umod* KI lines, we wanted to obtain a global view of the dynamic (mal)adaptive pathways operating in the mutant kidneys. RNA-sequencing (RNA-seq) of whole-kidney lysates revealed a strong clustering of the *Umod*^R186S/+^ samples, with a marked effect of age, in contrast with a closer proximity between the *Umod*^+/+^ and *Umod*^C171Y/+^ samples (<u>Supplementary Figures 6A, B</u>).

Compared to *Umod*^+/+^, the *Umod*^R186S/+^ kidneys showed an early (1 month) signature of ER stress response, increased inflammation and downregulation of lipid metabolic pathways (<u>Supplementary Figures 6C, D; Supplementary Table 7</u>), with a later (4 months) increase in immune response and activation of profibrotic pathways (<u>Supplementary Figures 7A, b</u>; <u>Supplementary Table 8</u>). Transcripts of TAL-enriched genes (*Umod, Car3, Egf*) were massively downregulated in the *Umod*^R186S/+^ kidneys. In contrast, the transcriptomic profile of *Umod*^C171Y/+^ kidneys was virtually indistinguishable from that of *Umod*^+/+^ kidneys at 1 month, with only subtle changes, e.g., in transcriptional regulation (*Jun, Btg2, Ier2, Fos*), observed at 4 months (<u>Supplementary Figure 7C-E; Supplementary Table 9</u>).

Comparison of the RNA profiles in *Umod*^R186S/+^ *vs. Umod*^C171Y/+^ kidneys gave insight into the differential adaptive and maladaptive response operating in the mutant kidneys. At 1 month (776 DEGs), activation of ER stress markers (*Trib3, Stc2, Nupr1*) and of the immune response (*Lcn2, Cxcl10, Lgals3*), with dysregulation of lipid metabolism (e.g., *Acat3, Acox2*) and ion transport (*Kcnt1, Clcnka, Slc5a3*), were noted in *Umod*^R186S/+^ *vs. Umod*^C171Y/+^ kidneys (<u>Supplementary Figures 8A, B</u>; <u>Supplementary Table 10</u>). At 4 months (1,461 DEGs), increased inflammation (*Lcn2, Dpt, Slc7a11*) and decreased TAL transcripts (*Umod, Egf, Kcnt1*) were observed in *Umod*^R186S/+^ *vs. Umod*^C171Y/+^ kidneys, whereas an enrichment of genes involved in transcriptional regulation (*Jun, Fos, Ier3*) or associated with protein folding (*Dnajb1, Hspb1, Cryab, Dnaja4, Hsp90ab1*) was observed in *Umod*^C171Y/+^ vs. *Umod*^R186S/+^ kidneys (Figures 6A, B; <u>Supplementary Table 11</u>). The activation of different ER stress response effectors, with prominent inflammation/fibrosis in *Umod*^R186S/+^ and protein folding enrichment in *Umod*^C171Y/+^ (Figure 6C, D), was validated by RT-qPCR analysis (Figure 6E).

**Figure 6.**
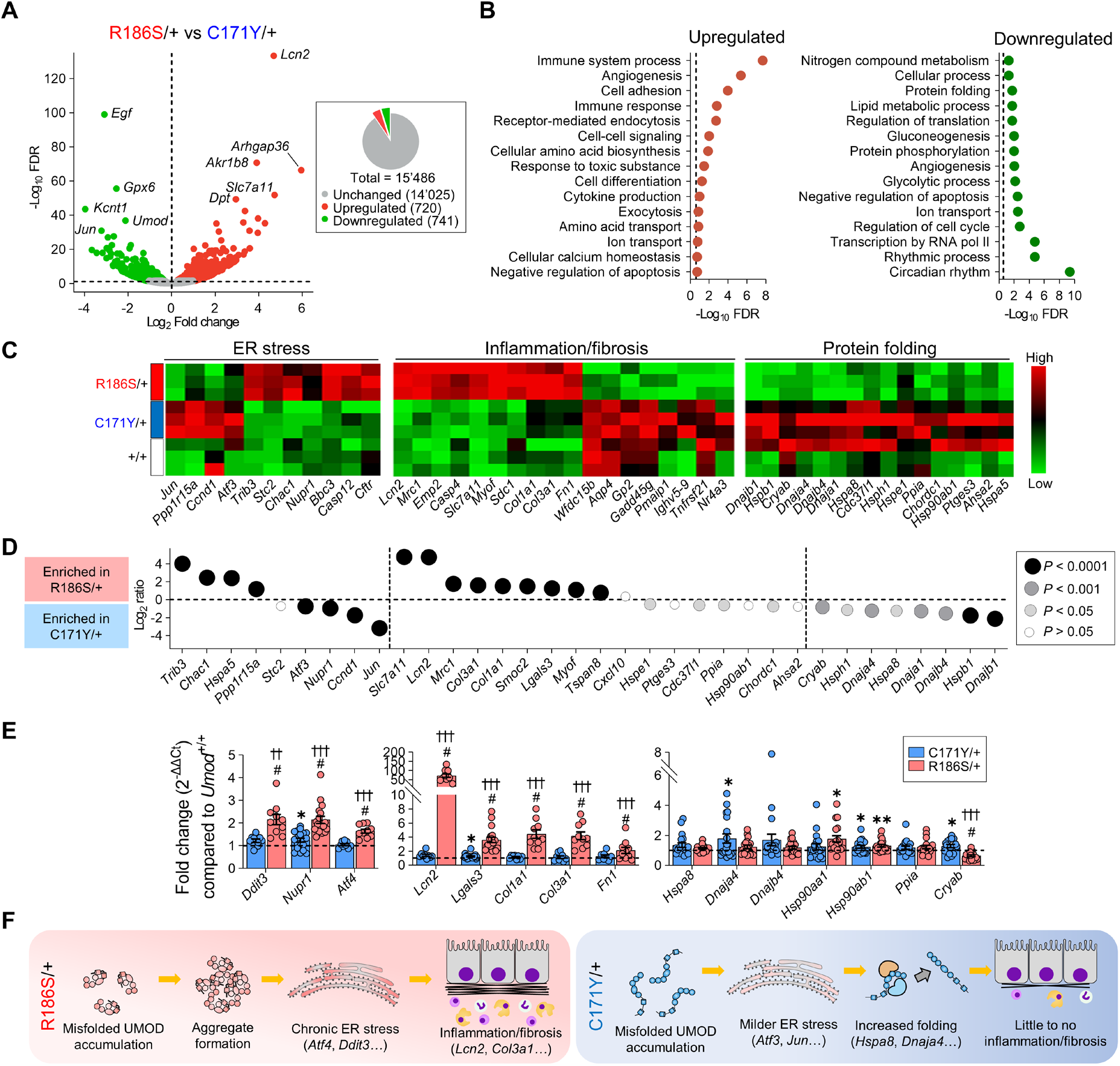
Distinct pathways impacting disease progression are activated in *Umod* KI kidneys. (**A**) Volcano plot showing differentially expressed genes (DEGs) between *Umod*^C171Y/+^ and *Umod*^R186S/+^ kidneys at 4 months. Genes not significantly changed (*FDR* > 0.05) are shown in gray, whereas genes that are up- or downregulated in *Umod*^R186S/+^ compared to *Umod*^C171Y/+^ are shown in red and green, respectively. The total numbers of unchanged, up- and downregulated genes are summarized in the pie chart. (**B**) Pathways of biological processes in gene ontology that are up- and downregulated in *Umod*^R186S/+^ compared to *Umod*^C171Y/+^. (**C**) Heat map of selected, divergent pathways involved in disease progression of *Umod* KI mice at 4 months. (**D**) Bubble plot of selected genes for key pathways, showing distinct signatures in the *Umod*^C171Y/+^ and *Umod*^R186S/+^ kidneys. (**E**) Validation of selected targets by RT-qPCR on 4-month kidneys (n= 9 to 19 animals per group). Values are relative to *Umod*^+/+^ (black dotted line). Bars indicate the mean ± SEM. One-way ANOVA with Tukey’s post hoc test; \**P* < 0.05, \*\**P* < 0.01, #*P* < 0.0001 versus *Umod*^+/+^; †*P* < 0.05, ††*P* < 0.01, †††*P* < 0.001 *Umod*^C171Y/+^ versus *Umod*^R186S/+^. (**F**) Schematic representation of pathophysiological mechanisms in *Umod*^R186S^ (left) and *Umod*^C171Y^ kidneys (right).

Together, these transcriptomic signatures indicate that *Umod*^R186S/+^ mice recapitulate typical/severe ADTKD-*UMOD*, with induction of ER stress response and development of inflammation and fibrosis. In contrast, *Umod*^C171Y/+^ kidneys show a milder and delayed activation of ER stress and increased expression of protein folding chaperones that presumably favor the processing and trafficking of uromodulin, resulting in considerably less inflammation and fibrosis (Figure 6F).

### *UMOD*-*GFP* cells recapitulate uromodulin trafficking defects and aggregates and activate specific quality control pathways

To gain mechanistic insight into mutant uromodulin processing, we transduced immortalized mouse inner medullary collecting duct (mIMCD-3) cells with GFP-tagged human wild-type (WT) or mutant (p.R185S, p.C170Y) *UMOD*. These cells (hereafter referred to as *UMOD*-*GFP* cells) recapitulated the main biochemical features of ADTKD-*UMOD*, including the formation of HMW (∼above 250 kDa) aggregates (Figure 7A) that were resolved by treatment with DTT (<u>Supplementary Figure 9A</u>), accumulation of EndoH-sensitive premature uromodulin (Figure 7B) and reduced uromodulin secretion (Figure 7C). Wild-type uromodulin is trafficked properly to the plasma membrane (Figure 7D), whereas both mutated isoforms show ER retention (Figure 7E).

**Figure 7.**
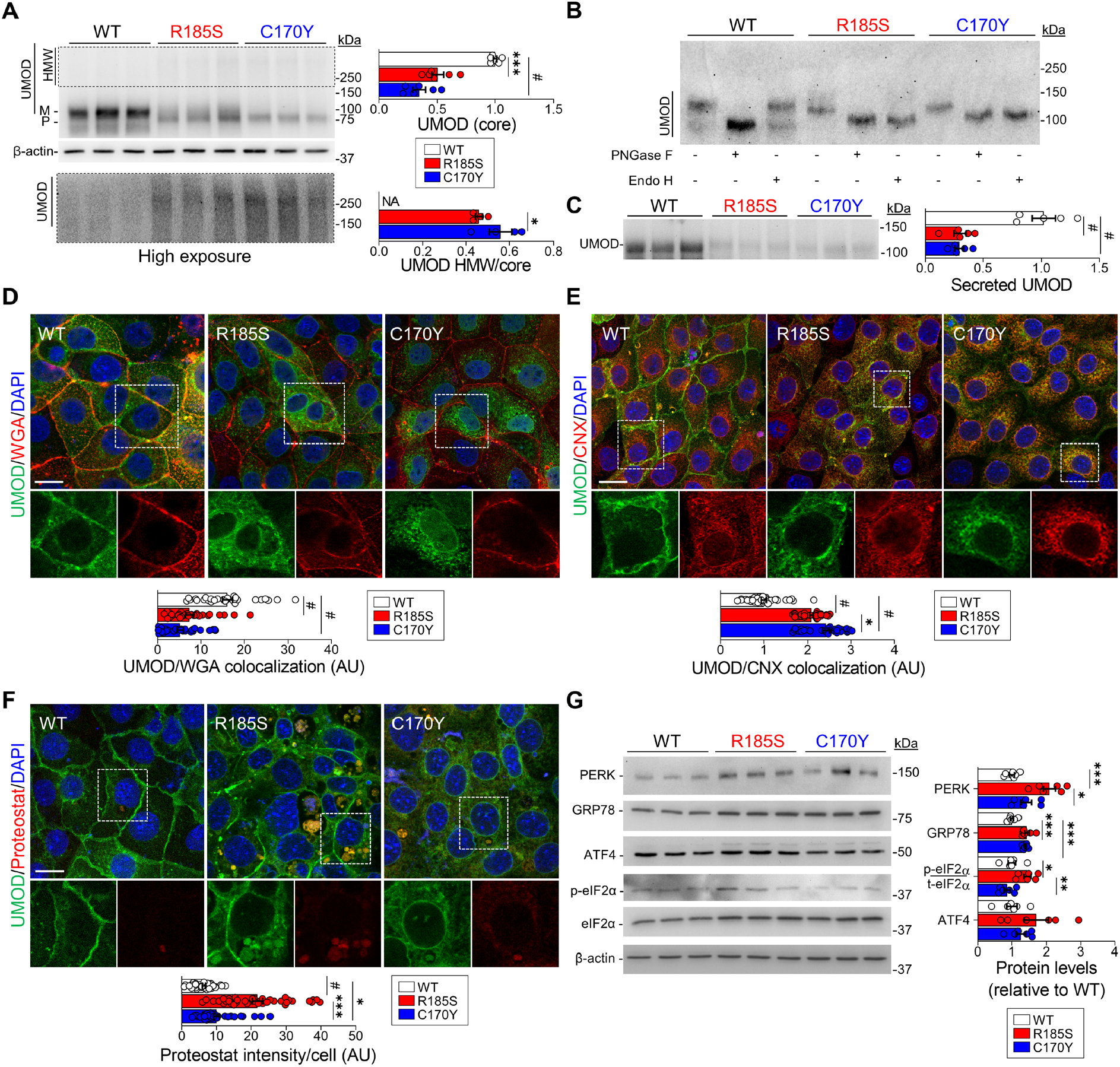
In vitro analysis of mutant uromodulin trafficking in mIMCD cells. (**A**) Immunoblot analysis of uromodulin (UMOD) in *UMOD*-*GFP* cell lysates. β-actin used as a loading control (*n* = 6 biological replicates). M: mature, P: precursor, HMW: high molecular weight. Bottom panel: longer exposure of HMW UMOD aggregates. (**B**) Immunoblot of UMOD from *UMOD*-*GFP* cell lysates following PNGaseF or EndoH treatment. (**C**) Representative immunoblot of UMOD in the apical medium of *UMOD*-*GFP* cells (*n*=5 biological replicates per group) (**D**-**F**) Representative immunofluorescence analysis of UMOD (green) and WGA (**D**), calnexin (CNX) (**E**) or PROTEOSTAT® (**F**) (red) in *UMOD*-GFP cells. White squares indicate images shown at higher magnification. Scale bar: 30 μm (**D, E**), 15 μm (**F**). (*n* = 29 to 39 cells per genotype. (**G**) Immunoblot analysis of UPR markers in *UMOD*-*GFP* cell lysates. β-actin used as a loading control (*n* = 6 to 9 biological replicates per group). Quantification is relative to UMOD WT. Bars indicate the mean ± SEM. One-way ANOVA with Tukey’s post hoc test, \**P* < 0.05, \*\**P* < 0.01, \*\*\**P* < 0.001, #*P* < 0.0001.

The localization of intracellular aggregates was investigated using the PROTEOSTAT^®^ dye, which labels aggresomes and related amyloid-like aggregates. Compared to WT cells, both mutant *UMOD* cell lines showed an enrichment for the PROTEOSTAT dye signal, colocalizing with uromodulin at the perinuclear area; the signal was significantly higher in R185S cells (Figure 7F). The R185S PROTEOSTAT-positive structures appeared to be surrounded by calnexin, suggesting that uromodulin aggregates are localized into ER inclusions (<u>Supplementary Figure 9B</u>).

Analysis of the level of ER stress showed an increased signal for GRP78/BiP in both R185S and C170Y mutant cells, colocalizing with uromodulin (<u>Supplementary Figure 9C</u>). Coimmunoprecipitation showed that both wild-type and mutant uromodulin interacted with GRP78 (<u>Supplementary Figure 9D</u>). However, higher expression of PERK, p-eIF2α, ATF4 and CHOP at either the mRNA (<u>Supplementary Figure 9E</u>) or protein (Figure 7G) levels was detected in R185S cells, whereas increased levels of the *Xbp1* transcript were observed in C170Y cells (<u>Supplementary Figure 9E</u>).

When checking for quality control mechanisms, we detected higher levels of *Dnaja4* in C170Y cells, as in *Umod*^C171Y/+^ kidneys, confirming the involvement of the protein folding machinery (<u>Supplementary Figure 9F</u>). The C170Y cells also showed higher levels of the translocon component *Sec62* and a significant upregulation of ER-phagy genes (*Rtn3, Retreg1, Sec62*), in contrast with the downregulation of *Ccpg1* in R185S cells (<u>Supplementary Figure 9F</u>). These results suggest that the UPR is activated through the PERK and IRE1 branches in mutant *UMOD* cell lines, with specific quality control mechanisms activated in C170Y cells, supporting our in vivo findings.

### Clearance of mutant uromodulin relies on distinct degradation mechanisms

To test whether the observed differences in trafficking, aggregate number and UPR induction reflect distinct clearance systems of mutant uromodulin, we investigated the ubiquitin-proteasome system (UPS) and autophagy-lysosomal degradation in the cells exposed to either proteasome (MG132) or autophagy (bafilomycin A1) inhibitors (Figure 8A). Following MG132 treatment, which increased the ubiquitin-binding protein SQSTM1 in all cells, uromodulin levels strongly increased in C170Y-expressing cells, whereas no significant change was detected in *UMOD* WT and R185S cells (Figure 8B). Unlike in *UMOD* R185S cells, higher SQSTM1 puncta colocalizing with uromodulin were initially present in C170Y mutant cells and further accumulated in MG132-treated cells (<u>Supplementary Figures 10A, B</u>). These data suggest that mutant cells respond differently to UPS inhibition, with C170Y mutant uromodulin being mainly targeted to this pathway.

**Figure 8.**
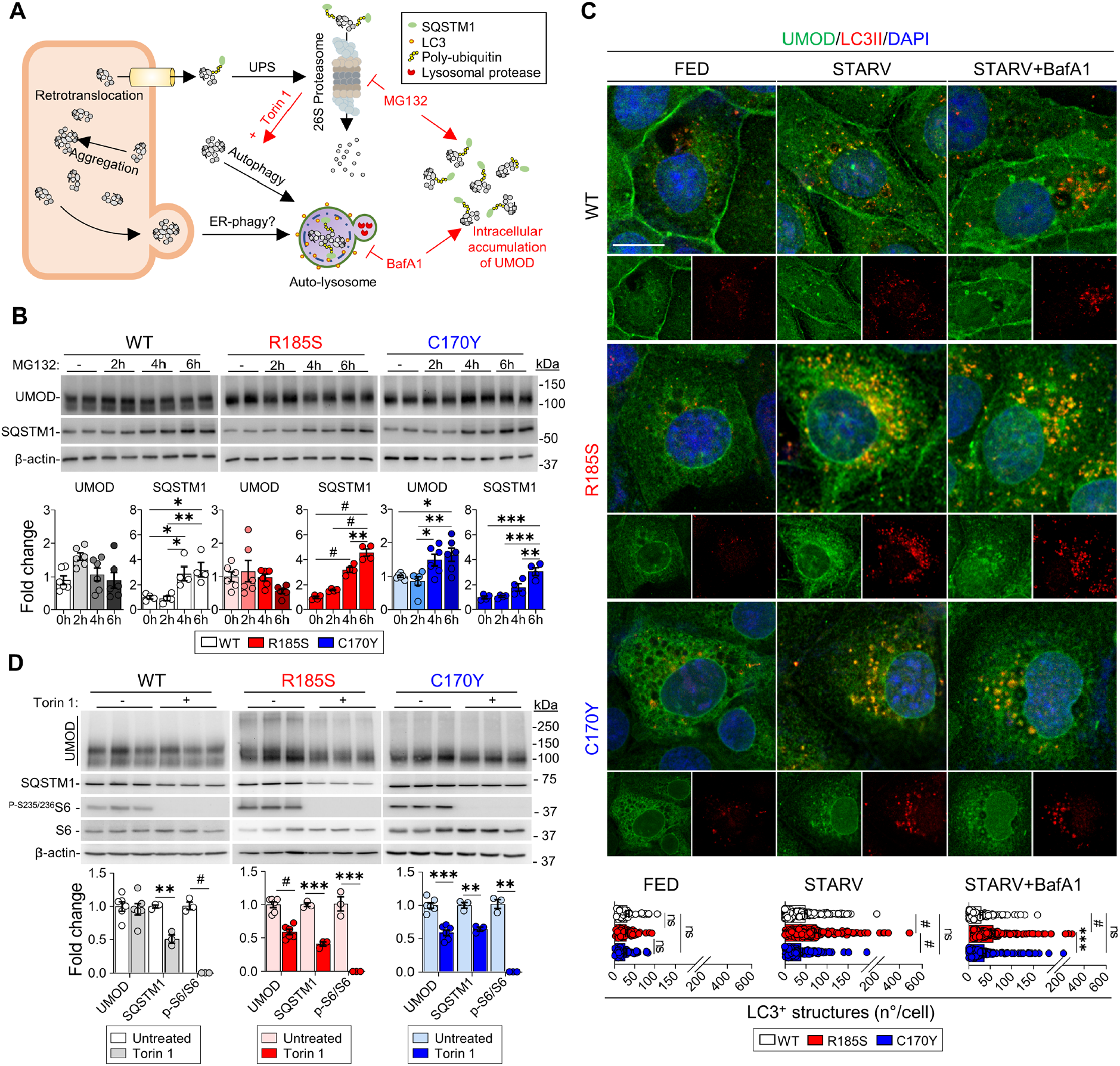
Degradative mechanisms involved in uromodulin clearance in vitro. (**A**) Diagram illustrating the degradation mechanisms of misfolded proteins and effect of different treatments. (**B**) Immunoblot analysis of UMOD and SQSTM1 in *UMOD*-*GFP* cells following MG132 treatment (*n* = 4 to 6 biological replicates per condition). (**C**) Immunofluorescence analysis of UMOD (green) and LC3 (red) in *UMOD*-*GFP* cells following starvation and bafilomycin A1 treatment. Nuclei are counterstained with DAPI (blue). Scale bar: 10 μm (*n* = 41 to 59 cells per condition). Separate channels are shown below each panel. (**D**) Immunoblot analysis of UMOD, SQSTM1 and mTOR effectors following treatment with Torin 1 in *UMOD*-*GFP* cells. β-actin was used as a loading control. Densitometry analysis relative to untreated samples. Bars indicate the mean ± SEM. One-way ANOVA with Tukey’s post hoc test, \**P* < 0.05, **, *P* < 0.01, \*\*\**P* < 0.001, #*P* < 0.0001.

Since R185S uromodulin did not increase following proteasomal inhibition, we investigated the potential involvement of the autophagy-lysosomal degradation machinery (Figure 8A). Starvation induced a significant decrease in uromodulin levels in R185S and C170Y cells, while levels in WT cells were unaffected (<u>Supplementary Figure 10C</u>). Treatment with BafA1 caused accumulation of uromodulin in R185S cells and C170Ycells, in contrast with no significant changes in WT cells (<u>Supplementary Figure 10C</u>). The increase in LC3II and SQSTM1 levels suggested no impairment of autophagic flux (<u>Supplementary Figure 10C</u>). After starvation, we detected a higher number of punctated LC3-positive structures colocalizing with uromodulin in R185S cells than in C170Y and WT cells, an effect that was further enhanced by BafA1 treatment (Figure 8C). Taken together, these data suggest that distinct uromodulin mutants are preferentially cleared by different mechanisms, with C170Y uromodulin being primarily degraded through the proteasome while R185S uromodulin is targeted to autophagy.

### Stimulating autophagy improves the clearance of mutant uromodulin

Considering the potential role of autophagy in the clearance of mutant uromodulin, we targeted this pathway by exposing the cells to the canonical mTOR complex 1 (mTORC1) inhibitor Torin 1. The efficient inhibition of mTORC1 signaling, as indicated by lower phosphorylation levels of mTORC1 target protein S6, was reflected by a significant decrease in both core and HMW uromodulin aggregates in the R185S and C170Y mutant cells (Figure 8D). Treatment with bafilomycin A1 (4 h following Torin 1) confirmed that the uromodulin reduction following Torin 1 treatment was due to autophagy induction. Bafilomycin A1 treatment inhibited lysosomal degradation and autophagy, as indicated by increased LC3B-II and SQSTM1 levels and significantly increased uromodulin levels in mutant cells compared to treatment with Torin 1 alone (<u>Supplementary Figure 10D</u>). Together, these data confirm the role of autophagy in the clearance of mutant uromodulin in the KI mouse models, opening therapeutic perspectives.

## DISCUSSION

ADTKD-*UMOD* is one of the most common monogenic kidney diseases, invariably causing kidney failure. A pressing research goal for ADTKD, as for other dominantly inherited diseases, is to bridge the gap linking postulated gain-of-function mutations with clinically relevant end points – here kidney fibrosis and progression to kidney failure. By combining human, mouse and cellular studies, we show how missense mutations of uromodulin, associated with divergent disease severity, lead to distinct trafficking defects and a propensity to form intracellular aggregates, with differential activation of ER stress and UPR, reflected by variable inflammation and end-organ damage. The *Umod*^C171Y/+^ mice showed a distinct protein folding chaperone response, preventing accumulation of premature uromodulin in the kidneys. Conversely, the *Umod*^R186S/+^ mice presented a massive accumulation of premature uromodulin in the kidney, with almost no residual uromodulin in the urine, triggering the formation of intracellular aggregates and a pathogenic cascade. Studies in kidney cells confirmed biochemical differences in the aggregates, with activation of distinct clearance mechanisms. Enhancement of autophagy by starvation and mTORC1 inhibition decreased uromodulin aggregates, suggesting a potential therapeutic strategy. These observations substantiate a model for allelic effects and the role of toxic aggregates in the progression of ADTKD-*UMOD*, with relevance for autosomal dominant disorders caused by toxic gain-of-function mechanisms.

More than 95% of the >135 *UMOD* mutations associated with ADTKD are missense, clustering in exon 3 and, in more than 50% of the cases, involving cysteine links (2). The *Umod* KI mouse models reported here aimed to investigate the impact of two mutations of *UMOD* (C170Y and R185S) associated with divergent clinical progression. Intuitively, one could assume that substitution of a conserved cysteine would negatively impact uromodulin stability through disruption of disulfide bridges, potentially triggering the formation of aggregates and disease progression (1, 2, 12). Previous studies comparing two *Umod* mutant mice obtained by ENU random mutagenesis concluded that the C93F mutation caused a more severe uromodulin maturation defect than A227T and a more severe phenotype (20). However, corresponding mutations to ENU mouse C93F and A227T are not reported in human ADTKD-*UMOD*. Furthermore, it was shown recently that among patients with ADTKD-*UMOD*, carriers of missense mutations involving a cysteine did not experience worse disease progression than those involving another residue (2). Analysis of clinical endpoints revealed that patients harboring the *UMOD* p. Arg185Ser mutation progressed to kidney failure much faster, by more than 25 years, than those with the p. Cys170Tyr mutation. This difference was reflected by findings on human kidney biopsies, where the p. Arg185Ser mutation was associated with ER accumulation, ER stress, and fibrosis, whereas the p. Cys170Tyr was predominantly detected on the apical plasma membrane, without induction of ER stress and no detectable fibrosis. Remarkably, these distinct disease progressions were substantiated in the two *Umod* KI lines, with strong allelic and gene-dosage effects on uromodulin processing and kidney damage and fibrosis.

While trafficking defects and accumulation of misfolded uromodulin in TAL cells have been reported in human kidney biopsies (12-14) and previous mutant *Umod* mouse models (18-22), little emphasis has been placed on the role of uromodulin aggregates. The availability of various KI lines, carefully matched for age and sex, allowed us to demonstrate the link between trafficking and maturation defects of uromodulin and the formation of HMW uromodulin aggregates in the kidney. These HMW aggregates, associated with the disappearance of mature uromodulin, were detected early and consistently in *Umod*^R186S/+^ kidneys, even at 2 weeks of age. In contrast, low levels of such aggregates were only detected in homozygous *Umod*^C171Y/C171Y^ kidneys, together with the mature form and ER-glycosylated uromodulin. Furthermore, the nature of the uromodulin mutation also influences the biochemical properties of the aggregates, whose formation involves both weak and strong interactions. The differences in the amount of biochemically distinct uromodulin were reflected in the differential induction of ER stress, as evidenced by the signal for GRP78/BiP and the PERK and IRE1α branches of the UPR in *Umod*^R186S^ but not in *Umod*^C171Y^ kidneys. Together, our quantitative analyses indicate close correlations between the amount of uromodulin aggregates, the induction of ER stress and UPR, the onset of inflammatory infiltrate and the extent of interstitial fibrosis in the kidneys of different *Umod* lines, substantiating the role of uromodulin aggregates in driving end-organ damage.

Importantly, deletion of the wild-type *Umod* allele in heterozygous *Umod*^R186S^ mice increased the amount of uromodulin aggregates and worsened inflammation, tubular damage markers and fibrosis. The fact that, despite the global reduction in uromodulin expression, there is increased aggregation, ER stress, and inflammatory parameters compared to the *Umod*^*R186S/+*^ kidneys strongly supports the pathogenic role of aggregates. Since *Umod* KO mice do not show any phenotype related to ADTKD-*UMOD* (27), these results suggest that wild-type uromodulin may protect against the formation of uromodulin aggregates. Studies on Huntington disease (HD), also characterized by a toxic gain of function, reported a similar protective role of wild-type huntingtin (Htt) protein in the disease: the absence of wild-type protein in HD mutant mouse and Drosophila models of Htt toxicity leads to a more severe phenotype (28, 29). The protective role of the wild-type allele may be to facilitate the trafficking of mutant protein to the plasma membrane or, alternatively, to directly balance the propensity of mutant uromodulin to form aggregates.

Our results suggest potential mechanisms accounting for the strong allelic and gene dosage effects on the formation of uromodulin aggregates and their toxicity to tubular cells. First, the transcriptional profiles revealed a canonical signature of ADTKD-*UMOD* progression, i.e. inflammation, interstitial fibrosis and kidney failure in the *Umod*^R186S^ kidneys, contrasting with an increased expression of genes involved in protein folding in the *Umod*^C171Y^ kidneys. The changes driven by the R186S mutation are in line with the emerging role of ER proteostasis in kidney disease, as evidenced for intermediate-effect size variants in *UMOD* (30) and for mutations in *MUC1* also associated with ADTKD (31). Furthermore, the strong induction of lipocalin-2 (LCN2) in the R186S kidneys may provide a link between ER stress, UPR, inflammation and fibrosis, as LCN2 is a known homing factor for inflammatory cells and an active player in kidney disease progression (32, 33). Recent structural data for human uromodulin show that the R185 residue falls into a very conserved region in the D10C domain that is required for the proper orientation of two critical N-glycans targeted by several mutations associated with ADTKD (34). Conversely, the increased folding capacity detected in the C171Y kidneys could explain why most of the uromodulin in these kidneys is terminally glycosylated and localized at the apical plasma membrane, with no decrease in its urinary excretion. Over time, these folding mechanisms may become metabolically challenging and/or insufficient to prevent the damage driven by the sustained production of mutant uromodulin (35).

Second, studies in R185S and C170Y cells demonstrated a different retention of mutant uromodulin altering ER homeostasis, reflected by differences in the formation of uromodulin aggregates – consistently higher in cells expressing R185S compared to the C170Y mutant. The upregulation of specific translocon components and ER-phagy genes suggests that specific quality control mechanisms are activated in C170Y cells, modulating the formation of toxic aggregates.

Third, investigations of UPS and autophagy-lysosomal degradation suggest that uromodulin mutants are differentially cleared: C170Y uromodulin is primarily degraded through the proteasome, while R185S uromodulin is targeted to autophagy. These results are in line with the fact that if chaperone-mediated folding cannot rescue misfolding, proteasomal degradation occurs since UPS is the main ER quality control mechanism (36). However, UPS can only degrade proteins, whereas ER components and protein aggregates are mainly cleared by autophagy-lysosomal degradation (37).

The central role of autophagy in clearing uromodulin aggregates offers perspectives for a therapeutic strategy. Previous studies suggested that autophagy could be impaired in ADTKD-*UMOD*, and rescued by mTOR inhibition (21). Here, induction of autophagy by either starvation or mTORC1 inhibition led to a decrease in mutant uromodulin and HMW aggregates in both mutant cell lines suggest the potential value of activators of autophagy in the clearance of mutant uromodulin. Further studies will be needed to validate this attractive possibility, taking advantage of experience in neurogenerative disorders and recently developed systems of targeted degradation of aggregation-prone proteins (38, 39).

In conclusion, we demonstrated that the differential effect of specific missense *UMOD* mutations on kidney disease progression is due to their propensity to misfold and aggregate, overcoming the capacity to activate chaperone and folding systems. The fact that toxic uromodulin aggregates drive end-organ disease implies that treatment strategies should aim to decrease uromodulin aggregation and/or to promote folding capacity. As in other dominant, toxic gain-of-function disorders, the protective role of wild-type uromodulin suggests the importance of allele-specific targeting to decrease the levels of uromodulin. In addition to activators of autophagy, activation of specific degradative pathways may be required to promote the clearance of specific uromodulin mutants.

## METHODS

### Human studies

For the assessment of kidney phenotypes according to *UMOD* mutations (age at onset of kidney failure and Kaplan-Meier analysis), only physician-verified data from ADTKD-*UMOD* patients/families recruited in Belgium or Switzerland were entered (in total, 78 patients from 30 families with 24 different *UMOD* mutations and with verified clinical data were analyzed). Kidney failure was defined as eGFR<15 ml/min/1.73 m^2^ or initiation of renal replacement therapy (dialysis or kidney transplantation). For a more detailed description of the study population, please refer to the respective figure legend.

### Animal experiments

Littermate animals of both sexes from three different colonies (*Umod*^R186S^, *Umod*^C171Y^ and *Umod*^R186S/KO^) were used all on the C57BL/6 background. The number of replicates for each experiment is indicated in the figure legends. Samples from the same experiment were processed simultaneously.

### Histological analysis and immunostaining

Immunodetection of uromodulin was performed on 5 μm-thick kidney sections obtained from nephrectomy samples of ADTKD-*UMOD* patients with p. Arg185Ser mutation in *UMOD* (female, 41 years old, ESKD; nephrectomy during kidney transplantation) and p.Cys170Tyr in *UMOD* (male, 58 years old, ESKD; nephrectomy due to kidney neoplasia). The slides were probed with sheep anti-uromodulin primary antibody, followed by Alexa Fluor 488-conjugated donkey anti-sheep. Coverslips were mounted with Prolong gold antifade reagent with 4′,6-diamidino-2-phenylindole (DAPI, P36931, Thermo Fisher Scientific) and analyzed under a Zeiss LSM 510 Meta Confocal microscope (Carl Zeiss, Jena, Germany) with high numerical aperture lenses (Plan-Neofluar 20x/0.5). Picrosirius Red staining (ab150681, Abcam, Cambridge, UK) was performed according to the manufacturer’s instructions. Briefly, samples were rehydrated and incubated for 1 hour in picrosirius red solution, rinsed briefly in 0.5% acetic acid, dehydrated in absolute ethanol, cleared and mounted. For mouse tissues, kidneys were harvested and processed as previously described (40,41). Five-micrometer-thick sections were incubated with the primary antibody for 2 hours at RT or overnight at 4 °C. The sections were then incubated with the appropriate AlexaFluor-conjugated secondary antibody (1:300, Life Technologies, Carlsbad, CA) for 1 hour at RT. The sections were mounted using Prolong Gold antifade reagent with DAPI (P36931, Thermo Fisher Scientific) and viewed under a confocal microscope (Leica Microsystems GmbH, Wetzlar, Germany) using a x63 1.4 NA oil immersion objective.

### Biochemical analysis and tissue collection

Urine samples were collected using individual metabolic cages (UNO Roestvastaal BV, Zevenaar, Netherlands) after appropriate training. Blood was collected by venous puncture or cardiac puncture at the time of sacrifice and centrifuged at 2000 g for 15 minutes at 4 °C in heparin-coated tubes (Sarstedt AG, Nürnbrecht, Germany) to separate plasma from cells. Creatinine, electrolytes, uric acid and urea were measured on a UniCel® DxC 800 Synchron® Clinical System or AU480 Clinical Chemistry System (Beckman Coulter, Brea, CA, USA) according to the manufacturer’s instructions. Urine and plasma osmolality were measured on a multisample osmometer (Advanced Instruments Model 2020, Norwood, MA, USA). Mice were anesthetized by intraperitoneal injection injection with a combination of ketamine (100 mg/ml) and xylazine (20 mg/ml) (Streuli Pharma AG, Uznach, Switzerland) in 0.9% NaCl for the collection of the kidneys. One kidney was flash frozen for RNA extraction, while the other was further processed for protein extraction and histological analyses. For medulla-enriched kidney fractions, kidneys were dissected in PBS under a Stereo microscope (VWR, SB350H VisiScope). Medulla- and cortex-enriched fractions were collected for protein extraction and validation of enrichment by Western blotting. For EM sample preparation, mice were anesthetized and perfused with a fixative solution containing 0.1 M sodium cacodylate (820670, Merck), 0.1 M sucrose (107687, Merck), 2% formaldehyde (#15710, Electron Microscopy Sciences, Hatfield, PA, USA) and 0.1% glutaraldehyde (#16220, Electron Microscopy Sciences). The kidneys were harvested, cut into ∼1 mm-thick slices and stored at 4 °C in fixative solution. Further processing was performed at the Center of Microscopy and Image Analysis of the University of Zurich.

### Cell models

mIMCD-3 (ATCC CRL-2123) cells were transduced with lentiviral particles for the expression of the indicated GFP-tagged human uromodulin isoforms (WT, C170Y and R185S). Lentiviral constructs were generated as described in Schaeffer et al. (42). The primers used for mutagenesis using the Quickchange Lightning mutagenesis kit (Stratagene, La Jolla, CA) are as follows:

hUmod_R185S_F 5’-GTACTCGGTGCTGCTCCAGTACTCGTCCA-3’;
hUmod_R185S_R 5’-TGGACGAGTACTGGAGCAGCACCGAGTAC-3’
hUmod_C170Y_F 5’-GGATCCGCGTACACGAGCGCGTCGC-3’;
hUmod_C170Y_R 5’-GCGACGCGCTCGTGTACGCGGATCC-3’.

### Protein sample preparation and immunoblotting

Kidneys and cells were lysed in radioimmunoprecipitation assay (RIPA) buffer (Sigma-Aldrich, St. Louis, MO, USA) containing protease and phosphatase inhibitors (Sigma-Aldrich) using an IKA T10 basic tissue homogenizer (IKA, Staufen, Germany) or cell scraper (99002, TPP Techno Plastic Products, Trasadingen, Switzerland). The lysates were briefly sonicated and centrifuged for 15 min at 1000 g and 4 °C to remove debris. Protein concentrations were determined using the bicinchoninic acid (BCA) protein assay kit (Thermo Fischer Scientific, Waltham, MA, USA), while urine was loaded according to creatinine concentration. For native PAGE, kidney lysates were diluted in sample buffer containing 0.01% bromophenol blue (Sigma-Aldrich), 25% glycerol and 12.5% Tris-HCl buffer, pH 6.8 (Bio-Rad, Hercules, CA, USA) and were run on 4-20% Mini-PROTEAN TGX Gel (Bio-Rad) in the absence of SDS. For deglycosylation analysis, samples were treated with PNGase F (P70404L, New England Biolabs, Ipswich, MA, USA) or Endo H (P0702, New England Biolabs) as previously described (41), according to the manufacturer’s instructions and analyzed by SDS-PAGE followed by immunoblotting. For semiquantitative immunoblot analysis, samples were reduced with DTT (except for uromodulin) and denatured by boiling. Samples were diluted in Laemmli sample buffer (Bio-Rad), separated on a 7.5–12% SDS-PAGE gel and blotted onto methanol-activated PVDF membranes. Membranes were blocked for 1 hour in 5% w/v nonfat dry milk solution, followed by overnight incubation at 4 °C with primary antibodies. Blots were subsequently incubated with peroxidase-conjugated secondary antibodies and visualized by Western HRP Substrate (Millipore, Burlington, MA, USA). Immunoblots were quantified by densitometric analysis using ImageJ or Image Lab.

### Solubility assays

Solubility assays on kidneys from *Umod* KI mice were based on a protocol to extract Sarkosyl-insoluble material from brain tissue (43). Minor modifications were implemented: starting material of 20 mg of frozen kidneys, resuspension of first insoluble pellets in 20 μl of buffer (20 mM Tris-HCl pH 7.4, 150 mM NaCl), no DTT treatment and resuspension of final pellets in 15 μl of 20 mM Tris-HCl pH 7.4, 150 mM with 30 seconds sonication. Ten micrograms of soluble material and total volume of insoluble material supplemented with Laemmli were boiled for 5 minutes and analyzed by Western blotting.

### Electron microscopy

For transmission electron microscopy (TEM), 1 mm thick kidney sections were immersed in 0.1 M cacodylate buffer containing 100 mM sucrose and 2.5% glutaraldehyde and embedded in epoxy resin. Correlative light-electron microscopy (CLEM) was performed as previously published (44). Briefly, sections were collected on wafers and washed with PBS (0.1 M, pH 7.4, 0 °C), followed by incubation with 0.15% glycine in PBS (3 × 1 min) and washing with PBS (3 × 1 min). Blocking solution composed of 0.5% BSA (albumin fraction V, Applichem) and 0.2% gelatin (from bovine skin, Sigma) in PBS was applied for 5 min, and samples were incubated with anti-uromodulin antibody (1:20) dissolved in blocking solution for 40 min at RT. After washing with blocking solution (6× 20 sec), the wafers were incubated with anti-sheep Alexa 647 secondary antibody (1:100, Life Technologies) in blocking solution (40 min at RT). After immunolabeling, sections were incubated with DAPI (4′,6-diamidino-2-phenylindole dihydrochloride, Roche, 1:250 dilution) for 10 sec and washed in PBS.

### RNA isolation, reverse transcription and quantitative PCR

Total RNA was extracted from kidney and cells using Aurum TM Total RNA Fatty and Fibrous Tissue Kit (Bio-Rad) or RNAqueous™ Micro Total RNA Isolation Kit (Thermo Fisher Scientific) following the manufacturer’s protocol. Analysis of gene expression levels in mouse kidneys and cells was performed based on the MIQE guidelines. Reverse transcriptase reaction with iScript TM cDNA Synthesis Kit (Bio-Rad) was executed with up to 1 μg of RNA. When needed, PCR products were size fractionated on a 1.5% agarose gel and stained with EZ-VisionR One (AMRESCO, Solon, OH). The variations in mRNA levels of the target genes were established by relative RT-qPCR with a CFX96TM Real-Time PCR Detection System and the iQ™ SYBR Green Supermix (Bio-Rad) for the detection of single PCR product accumulation. Primers specific to targets were designed with Beacon Design 2.0 (Premier Biosoft International, Palo Alto, CA). The PCR conditions were: 95 °C for 3 min, followed by 40 cycles of 15 seconds at 95 °C and 30 seconds at 60 °C. The efficiency of each set of primers was determined by dilution curves (<u>Supplementary Table 12</u>). GeNorm version 3.4 was used to characterize the expression stability of the candidate reference genes in the kidney. The normalization factor was determined using 6 housekeeping genes (*18S, 36B4, Actb, Gapdh, Hprt1*, and *Ppia*) for tissue samples. The relative changes were determined by the formula: 2^−ΔΔct^ and expressed as a percentage relative to *Umod*^+/+^ mice or *UMOD* WT cells.

### RNA-sequencing and bioinformatic analysis

RNA sequencing was performed at Eurofins Genomics (Ebersberg, Germany) using an Illumina HiSeq 2500 sequencer (Illumina, San Diego, CA, USA). Information on RNA quality and yield can be found in the Supplementary Data (<u>Supplementary Table 13</u>). Network Analyst 3.0 (45) was used for processing and analysis of RNA-Seq data, principal component analysis (PCA), generation of heat maps and overrepresentation analysis (ORA). Data were log_2_-normalized and analyzed using the DESeq2 algorithm (46). Volcano plots and ORA scatterplots were generated with GraphPad Prism. Primary RNA-sequencing data are available on GEO (accession number: GSE214491).

### Isolation of TAL segments from mouse kidneys

The TAL segments were isolated from mice as previously described (47). Briefly, kidneys were harvested, the outer medulla was cut into 1-mm-square pieces, digested for 30 minutes with 245 units/ml type-2 collagenase (Worthington Biochemical Corp, Lakewood, USA) and 96 μg/ml soybean trypsin inhibitor (Sigma), and segments were sieved through a 250-μm filter (BVBA Prosep, Zaventem, Belgium) and collected on an 80-μm filter (BVBA). TALs were selected under a light microscope based on morphology (Leica DMIL, Bensheim, Germany). Pooled TAL segments (∼100) were resuspended in 50 μL of Laemmli sample buffer (Bio-Rad) and analyzed by Western blot.

### Cell culture and treatments

For UPS treatment, *UMOD*-*GFP* cells were treated for 6 h with 2.5 μM proteasome inhibitor MG132 (M8699, Sigma-Aldrich). Protein samples were collected at 2, 4, and 6 hours post treatment for Western blot analysis. For the autophagy experiment, cells were incubated for 6 hours with serum-free medium (12440-061, IMDM Gibco) and for 4 hours with 250 nM bafilomycin A1 (b1793, Sigma-Aldrich). For mTOR treatment, cells were incubated for 16 h with 250 nM mTOR inhibitor Torin 1 (4247, TOCRIS Bioscience, Bristol, UK) and 4 h with 200 nM bafilomycin A1. Samples were analyzed by both Western blot and immunofluorescence analyses.

### Cells Immunofluorescence and image analysis

*UMOD*-*GFP* cells were fixed in 4% PFA, quenched with 50 mM NH_4_Cl, and permeabilized in blocking buffer solution containing 0.5% saponin and 0.5% BSA dissolved in PBS. Subsequently, cells were incubated overnight at 4 °C with the appropriate primary antibody or stained for 30 min at RT° with PROTEOSTAT^®^ dye (ENZ-51035-K100) as previously described (48). For the PROTEOSTAT^®^ Aggresome detection kit, all components were prepared according to the manufacturer’s instructions. The samples were incubated for 45 min with suitable fluorophore-conjugated Alexa secondary antibodies (1:400, Thermo Fisher Scientific), counterstained with DAPI (Invitrogen), mounted with Prolong Gold antifade reagent (P36931, Thermo Fisher Scientific) and analyzed by a Leica SP8 confocal laser scanning microscope (Leica Microsystems GmbH, Wetzlar, Germany). Quantitative image analysis was performed by randomly selecting five visual fields pooled from biological triplicates, with each field including at least 20–25 cells. The quantitative analyses were performed using open-source cell image analysis software for cell phenotype (CellProfiler) and ImageJ.

### Statistics

All values are expressed as the mean ± SEM. Prior to statistical analysis, sample populations were tested for normality (Shapiro-Wilk test) and equality of variances (*F*-test). Two-group comparisons were performed using either unpaired, two-tailed Student’s t-test or the Mann-Whitney test in cases of nonnormal distribution of the samples. Comparisons between three or more groups were performed using one-way ANOVA followed by Tukey-Kramer’s post hoc analysis. *P* values < 0.05 were considered statistically significant.

### Study approval

Clinical and genetic information for ADTKD-*UMOD* patients were retrieved from the Belgo-Swiss ADTKD registry, comprising a subset of patients recruited in the International ADTKD Cohort (2). Informed and written consent was obtained from all participants. The ADTKD registry was approved by the Institutional Review Board of the UCLouvain Medical School and Saint Luc University Hospital, Belgium (2011/04MAI/184) and the European Community’s Seventh Framework Programme European Consortium for High-Throughput Research in Rare Kidney Diseases (EURenOmics) Ethics Advisory Board and its participating centres. The use of human samples was approved by the UCLouvain Ethical Review Board. All animal experiments were performed in accordance with the ethical guidelines at University of Zurich (Zurich, Switzerland) and the legislation of animal care and experimentation of Canton Zurich, Switzerland. The experimental protocols were approved by the appropriate licensing committee (Kanton Zürich Gesundheitsdirektion Veterinäramt; l ZH049/17 and ZH195/20) at the University of Zurich.

## Author contributions

G.S., J.L., M.M., L.R., E.O. and O.D. conceptualized the study. O.D. supervised the work. G.S., J.L. and M.M. performed the in vivo and derived studies, providing the basis of the co-first authorship order. M.H. performed in vivo and in vitro experiments. E.O. provided human genetic data. C.S. and L.R. provided the *UMOD*-*GFP* cells. G.S., J.L., M.M. and E.O. analyzed and interpreted the data. G.S. performed the RNA-seq analysis. G.S., J.L., M.M., E.O. and O.D. wrote the paper with inputs and comments from all the authors. All authors reviewed the final version of the manuscript.

## Acknowledgments

E.O. is supported by the Swiss National Science Foundation (P2ZHP3_195181 and P500PB_206851) and Kidney Research UK (Paed_RP_001_20180925). M.M and O.D. are supported by the European Union’s Horizon 2020 research and innovation program under the Marie Skłodowska-Curie grant (agreement N° 860977). O.D. is supported by the European Reference Network for Rare Kidney Diseases (project N° 739532), the Swiss National Science Foundation’s National Center of Competence in Research Kidney Control of Homeostasis program, the Swiss National Science Foundation (grant 310030-189044), and the University Research Priority Program (URPP) ITINERARE at the University of Zurich. L.R. is supported by the Italian Ministry of Health (RF-2010-2319394 and RF-2016-02362623).

We thank Aleksandra Kokanovic, Nadine Nägele and Huguette Debaix for expert technical assistance, Ines Dufour and Nathalie Demoulin for clinical information on ADTKD families, Selda Aydin for material, Merel van Gogh for assistance with cell sorting and Jessica J. Stanisich and Rudi Glockshuber (ETH Zürich) for fruitful scientific discussion.

## Notes

The authors have declared that no conflict of interest exists.

### Competing Interest Statement

The authors have declared no competing interest.

### Summary of Updates

Changes have been implemented in the abstract, main text, figures and supplementary data. Quality of the figures has been improved.

