## Supplementary Materials for "Allelic and Gene Dosage Effects Involving Uromodulin Aggregates Drive Autosomal Dominant Tubulointerstitial Kidney Disease"

### Supplementary Material

Supplementary Methods
Supplementary Figure 1: Uromodulin characteristics in the C171Y and R186S mouse models.
Supplementary Figure 2: Early uromodulin processing defects in R186S/+ mice.
Supplementary Figure 3: Unfolded protein response in kidneys from *Umod* KI mice.
Supplementary Figure 4: Lack of apoptosis or caspase activation in *Umod* KI kidneys.
Supplementary Figure 5: Mutant UMOD degradation relies on mutation-specific mechanisms.
Supplementary Figure 6: RNA-seq study design, differential expression and affected pathways in 1-month-old R186S/+ kidneys.
Supplementary Figure 7: Differential expression and affected pathways in 4-months-old *Umod* KI kidneys.
Supplementary Figure 8: Disease progression signature in *Umod* KI kidneys.
Supplementary Figure 9: Distinct UMOD mutations trigger differential ER quality control responses.
Supplementary Figure 10: Autophagy induction enhances mutant UMOD clearance.
Supplementary Table 1: Clinical characteristics of p. (Arg185Ser) ADTKD-*UMOD* family.
Supplementary Table 2: Clinical characteristics of p. (Cys170Tyr) ADTKD-*UMOD* families.
Supplementary Table 3: *In silico* analysis of selected *UMOD* missense variants.
Supplementary Table 4: Clinical and biochemical parameters of *Umod*$^{C171Y}$ mice.
Supplementary Table 5: Clinical and biochemical parameters of *Umod*$^{R186S}$ mice.
Supplementary Table 6: Clinical and biochemical parameters of *Umod*$^{R186S/-}$ mice.
Supplementary Table 7: Top 50 DEGs in *Umod*$^{R186S/+}$ kidneys at 1 month.
Supplementary Table 8: Top 50 DEGs in *Umod*$^{R186S/+}$ kidneys at 4 months.
Supplementary Table 9: Top 50 DEGs in *Umod*$^{C171Y/+}$ kidneys at 4 months.
Supplementary Table 10: Top 50 DEGs in *Umod*$^{R186S/+}$ compared to *Umod*$^{C171Y/+}$ kidneys at 1 month.
Supplementary Table 11: Top 50 DEGs in *Umod*$^{R186S/+}$ compared to *Umod*$^{C171Y/+}$ at 4 months.
Supplementary Table 12: Primers used for real-time RT-PCR analysis.
Supplementary Table 13: RNA-Seq quality and yield.
Supplementary References

### Supplementary Methods

#### Generation of the *Umod* targeting vectors

Both targeting vectors contained 8.8 kb of the *Umod* gene with the respective mutations, a FRT-flanked neomycin resistance cassette and a SalI restriction site that was used for linearization of the vectors prior to electroporation of C57BL/6N-derived embryonic stem (ES) cells. A total of 300 p.R186S and 400 p.C171Y clones were resistant to selection with Geneticin, with 23 p.R186S and 24 p.C171Y clones that had undergone homologous recombination and were subsequently expanded. Presence of the mutations was established by restriction/PCR combined analysis and by Southern blot (data not shown). Selected positive clones were injected in blastocysts from grey C57BL/6 mice (R186S: 4 clones in 60 blastocysts; C171Y: 2 clones in 36 blastocysts) and the surviving blastocysts were transferred in CD-1 foster mice. The resulting highly chimeric mice (80-100%) were mated to grey C57BL/6N Flp-deleter mice to obtain offspring in which the neomycin resistance cassette was excised.

#### *In situ* cell death detection

For apoptosis labeling, Terminal transferase dUTP nick end labeling (TUNEL) assay was performed with In Situ Cell Death Detection Kit, Fluorescein (11 684 795 910, Roche, Basel, Switzerland), according to the manufacturer's instructions. The labeling procedure for difficult tissue was used. Briefly, slides were deparaffinized in xylene and rehydrated in a graded ethanol series. Antigen retrieval was carried out for 10 minutes with citrate buffer (pH 6.0) at 98°C, following by blocking for 30 minutes in Tris-HCl 0.1M (pH 7.4) containing 3% BSA and 20% normal bovine serum. Slides were incubated with the TUNEL reaction mix for 1 hour at 37 °C. Slides were then probed with sheep anti-uromodulin primary antibody (1/400; Meridien Life Science Inc. K90071C) for 1 hour at room temperature, followed by AlexaFluor633-conjugated donkey anti-sheep (1/400; Invitrogen). Coverslips were mounted with Prolong gold antifade reagent with DAPI (P36931, Thermo Fisher Scientific) and analyzed under a Confocal microscope (Leica Microsystems GmbH) using a ×63 1.4 NA oil immersion objective.

#### Co-immunoprecipitation

*UMOD-GFP* cells were grown to confluence in T75 flasks and lysed in 1 mL of RIPA buffer containing protease and phosphatase inhibitors, for 15 min at 4°C followed by 10 min centrifugation at 15,000g. Cell lysates (1 mg) were incubated under rotation overnight at 4°C with 50 µl protein G-Sepharose beads (pre-conjugated with 3 µl of rabbit anti GRP78 antibody (ab21685, Abcam) or with 6 µl of sheep anti-uromodulin antibody (K90071C, Meridian Life

Science Inc., Cincinnati, OH, USA). Beads were washed 3 times in PBS and the immunoprecipitated material was eluted with 20 µl of 50 mM glycine pH 2.8 and 1M Tris-HCl pH 7.4 was used to neutralize the samples. Samples were boiled 5 min at 95°C for Western Blot analysis.

**Semiquantitative analysis of fibrosis**

Full section-scans of 4 months-old *Umod*$^{+/+}$, Umod$^{R186S}$ and Umod$^{C171Y}$ kidneys were acquired at a resolution of 200× by a Zeiss Axio Scan.z1 slide scanner (Carl Zeiss, Oberkochen, Germany), equipped with a Hitachi HV-F202FCL digital video camera (Hitachi, Tokyo, Japan). Images were acquired using ZENblue software (Carl Zeiss SpA) and analyzed using ImageJ. Large blood vessels and inner medulla were manually excluded from quantification. Images were deconvoluted to obtain separate collagen signal (red) and cytoplasmic signal (yellow). Fibrotic area was calculated as percentage of the collagen area over total tissue area (collagen + cytoplasmic).

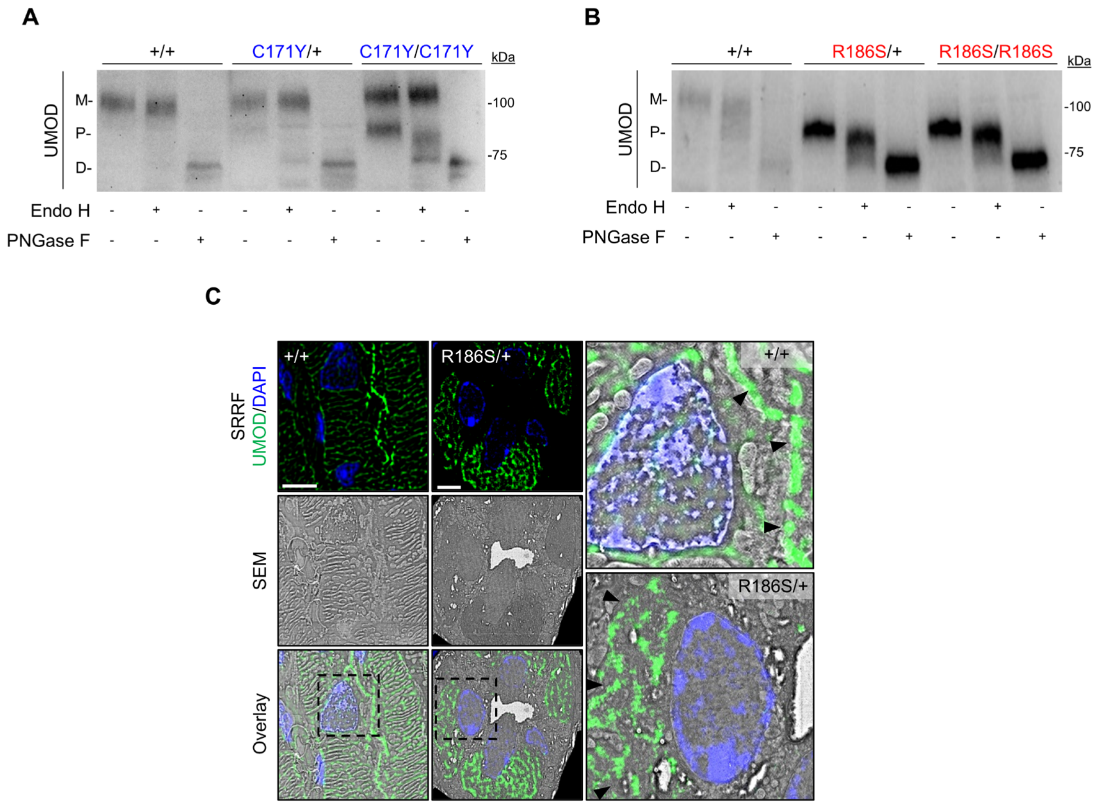

**Supplementary Figure 1: Uromodulin characteristics in the C171Y and R186S mouse models.**
(**A-B**) Immunoblot analysis of UMOD following Endo H or PNGase F treatment in kidneys from 1-month-old C171Y (**A**) and R186S (**B**) mice. M: mature; P: precursor; D: deglycosylated (**C**) Correlative Light Electron Microscopy (CLEM) of UMOD (green) on kidney sections from 3-month-old +/+ and R186S/+ mice. Nuclei are counterstained with DAPI (blue). Black arrowheads indicate UMOD localization (apical plasma membrane in +/+, endoplasmic reticulum in R186S/+). Scale bar: 5 µm. SRRF, super-resolution radial fluctuation; SEM, scanning electron microscopy.

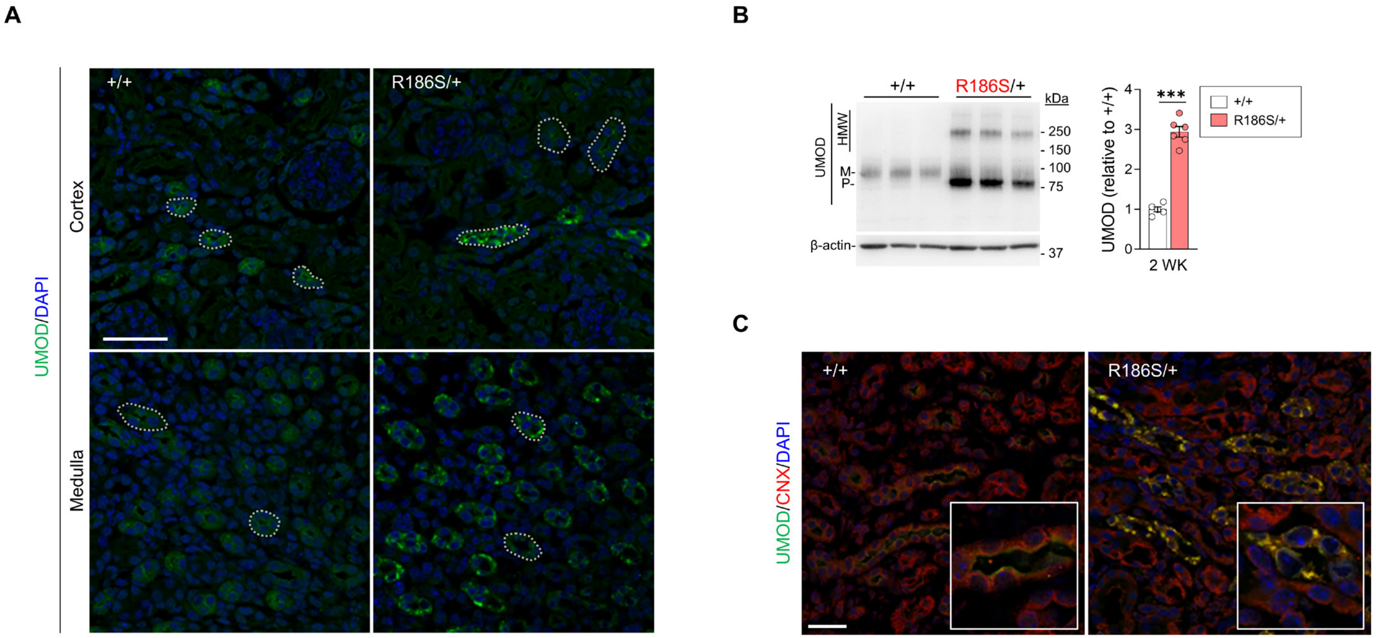

**Supplementary Figure 2: Early uromodulin processing defects in R186S/+ mice.**

(**A**) Representative immunofluorescence analysis of UMOD (green) on kidney sections from 2-week-old +/+ and R186S/+ mice. Nuclei counterstained with DAPI (blue). Scale bar: 25 µm. Dotted line identifies different tubules with apical or intracellular UMOD accumulation. (**B**) Representative immunoblot of UMOD in whole kidney samples from 2-week-old +/+ and R186S/+ mice. β-actin used as loading control. M: mature UMOD; P: premature UMOD; HMW: high molecular weight. Densitometry analysis relative to +/+. Bars indicate mean ± SEM. Unpaired two-tailed t test (right), ***$P < 0.001$ (n = 5 to 6 animals per group). (**C**) Representative immunofluorescence analysis of UMOD (green) and CNX (red) on kidney sections from 2-week-old mice. Nuclei are counterstained with DAPI (blue). Scale bar: 25 µm.

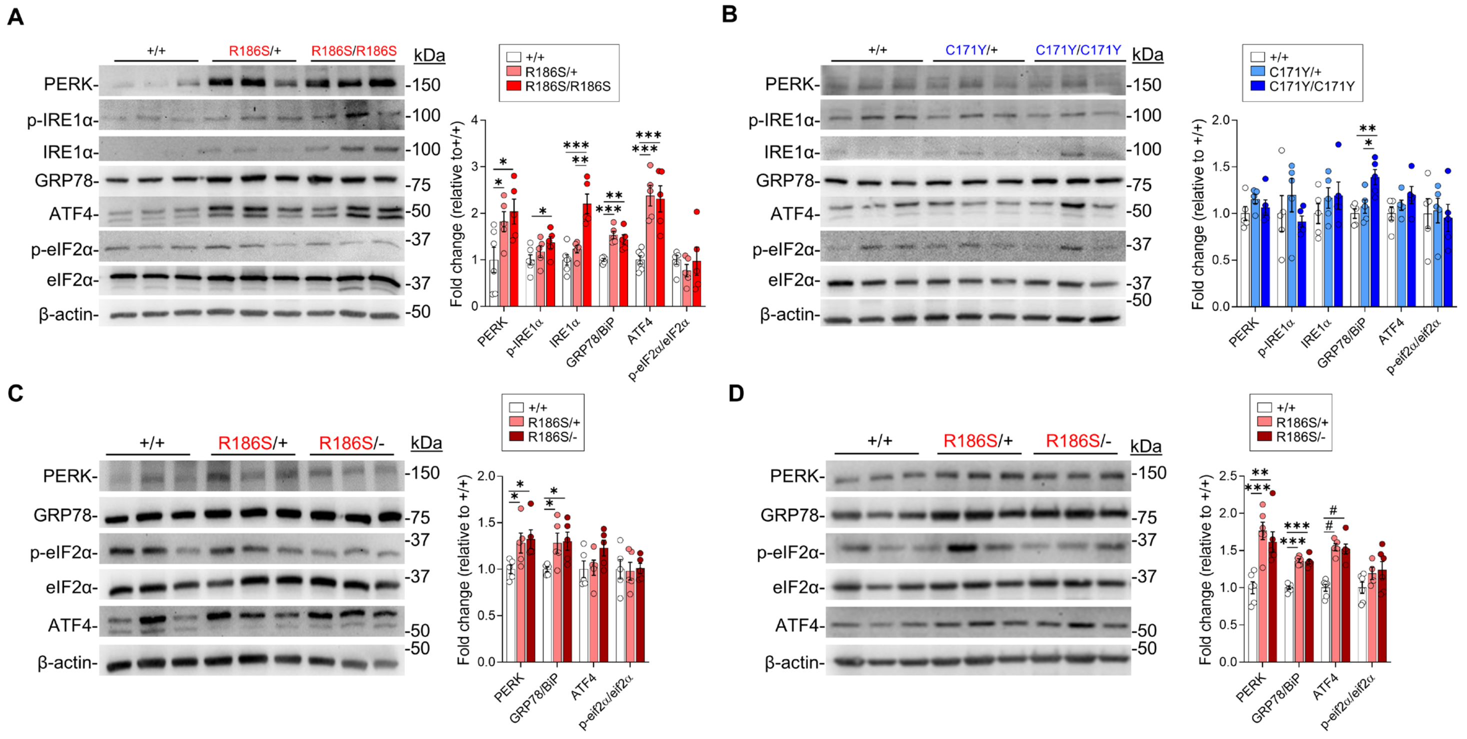

**Supplementary Figure 3: Unfolded protein response in kidneys from *Umod* KI mice.**

(**A-B**) Representative immunoblot analysis of ER stress markers in medulla-enriched kidney fractions from 4-months-old R186S (**A**) and C171Y (**B**) mice. (**C-D**) Representative immunoblot analysis of ER stress markers in 1-month-old medulla-enriched kidney fractions (**C**) or 4-months-old total kidney lysates from +/+, R186S/+ and R186S/- mice. β-actin used as loading control. Densitometry analysis is relative to +/+. Bars indicate mean ± SEM. One-way ANOVA followed by Tukey's post-hoc test, **P* < 0.05, ***P* < 0.01, ****P* < 0.001, #*P* < 0.0001 (n= 5 to 6 animals per group).

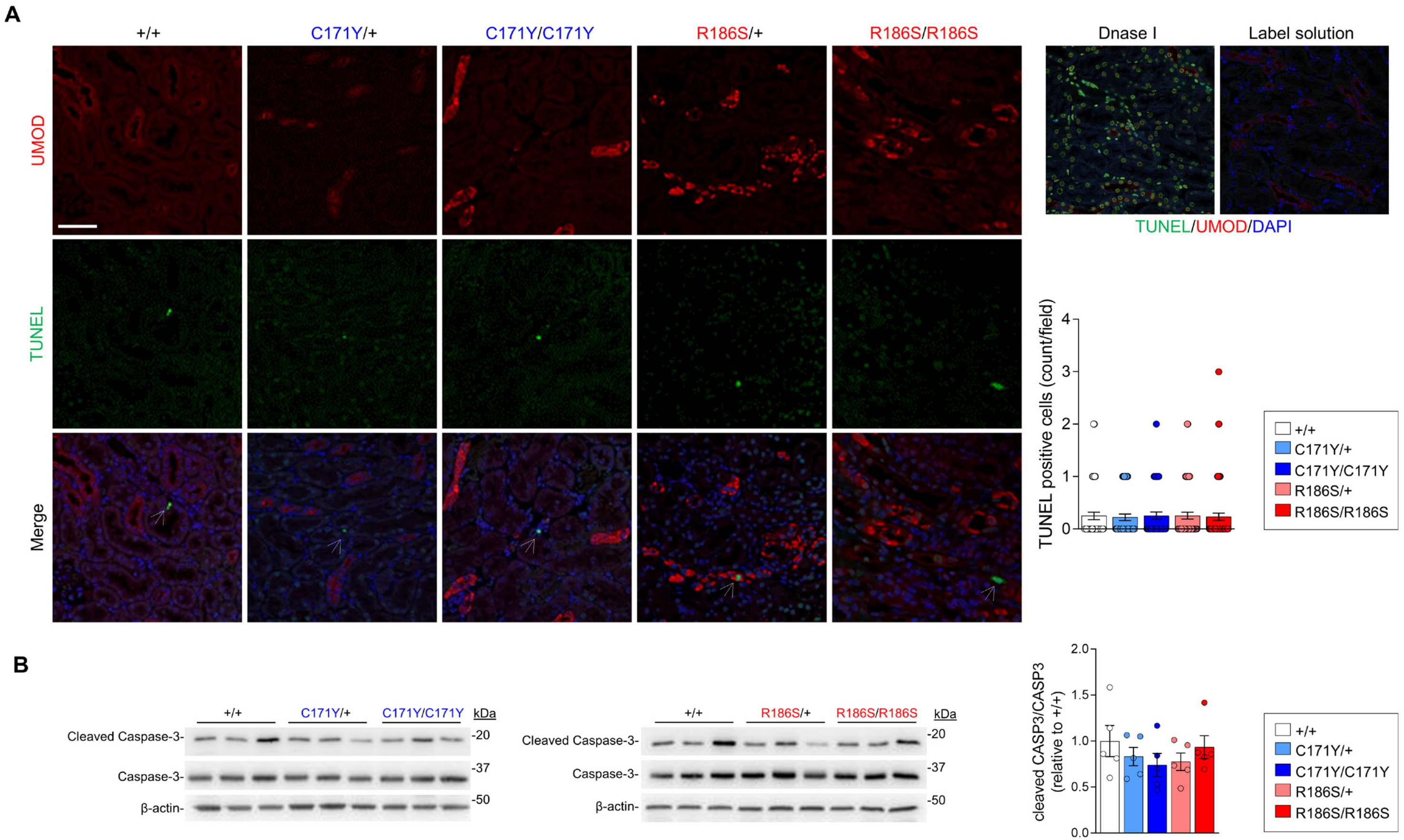

**Supplementary Figure 4: Lack of apoptosis or caspase activation in *Umod* KI kidneys.**
(**A**) Immunofluorescence analysis of TUNEL (green) and UMOD (red) in kidney sections from 4-month-old *Umod* mutant. Nuclei are counterstained with DAPI (blue). Positive control incubated with DNase I and negative control incubated with label solution shown on right panel. Scale bar: 50 µm. Each point of the quantification represents the number of TUNEL+ cells in one field. Bars indicate mean ± SEM. One-way ANOVA followed by Tukey's post-hoc test, n ≥ 45 fields from 3-5 kidneys per condition. (**B**) Representative immunoblot analysis of caspase-3 (CASP3) and cleaved caspase-3 in whole kidney lysates from 4-month-old +/+, C171Y/+ and C171Y/C171Y mice (left) or +/+, R186S/+ and R186S/R186S mice (right). β-actin used as a loading control. Densitometry analysis is relative to +/+. Bars indicate mean ± SEM. One-way ANOVA followed by Tukey's post-hoc test, (n = 5 animals per group).

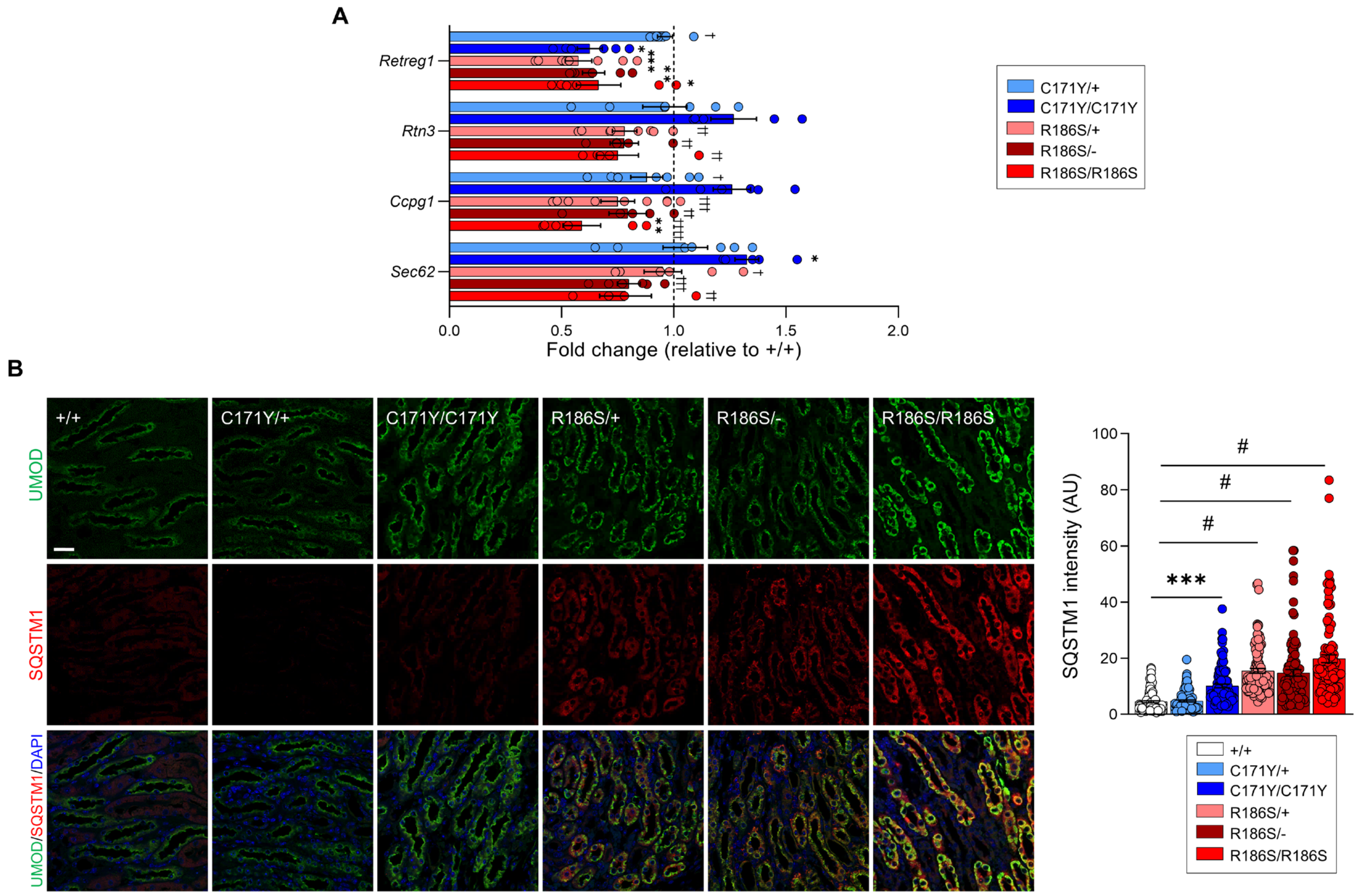

**Supplementary Figure 5: Mutant UMOD degradation relies on mutation-specific mechanisms.**

(**A**) RT-qPCR analysis of ER-phagy genes in *Umod* KI kidneys. Values are expressed as relative to +/+ (black dotted line), (n=4 to 9 animals per group). (**B**) Representative immunofluorescence analysis of UMOD (green) and SQSTM1 (red) on kidney sections from 1-month-old *Umod* KI mice. Nuclei are counterstained with DAPI (blue). Scale bar: 25 µm, n = 100 tubules from 3 kidneys per condition. Bars indicate mean ± SEM. One-way ANOVA followed by Tukey's post-hoc test, **P* <0.05, ***P* <0.01 ****P* < 0.001, #*P* < 0.0001, † *P* < 0.05, †† *P* < 0.01, ††† *P* < 0.001, †††† *P* < 0.0001. *compared to +/+, †compared to C171Y/C171Y.

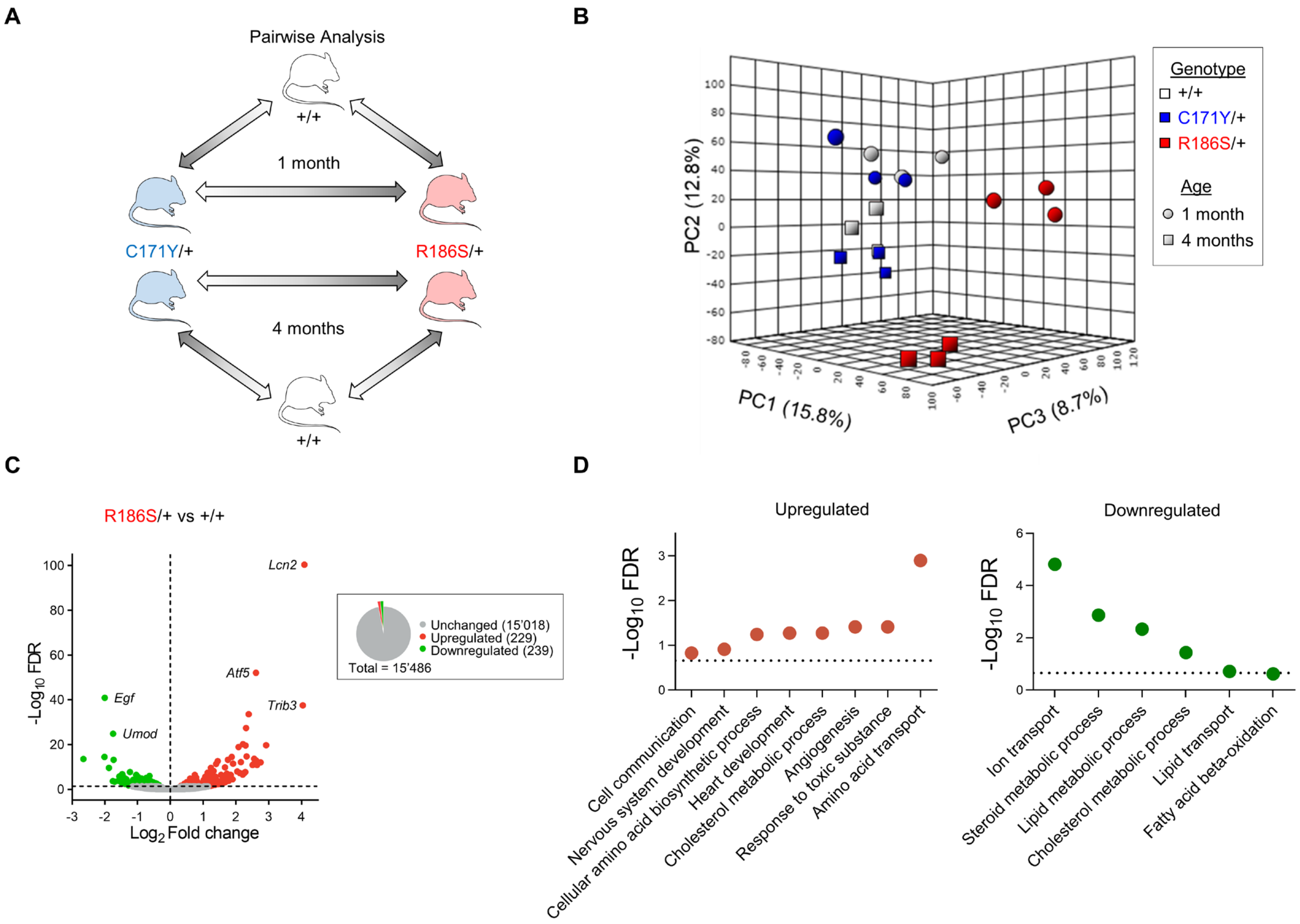

**Supplementary Figure 6: RNA-seq study design, differential expression and affected pathways in 1-month-old R186S/+ kidneys.**
(**A**) Experimental design for RNA-Seq on whole kidney lysates from 1 and 4 months *Umod* KI mice. (**B**) Principal component analysis (PCA) of RNA-Seq data of kidneys from 1-month-old and 4-month-old *Umod* KI mice. (**C**) Volcano plot showing differently expressed genes (DEGs) between R186S/+ and +/+ 1-month-old kidneys. Genes not significantly changed (FDR > 0.05) are shown in grey, whereas genes that are up- or downregulated in R186S/+ are shown in red and green respectively. The total number of unchanged, up- and downregulated genes are summarized in the pie chart. (**D**) Over-representation analysis showing up- (red) and downregulated (green) biological processes of gene ontology in 1-month-old R186S/+ kidneys.

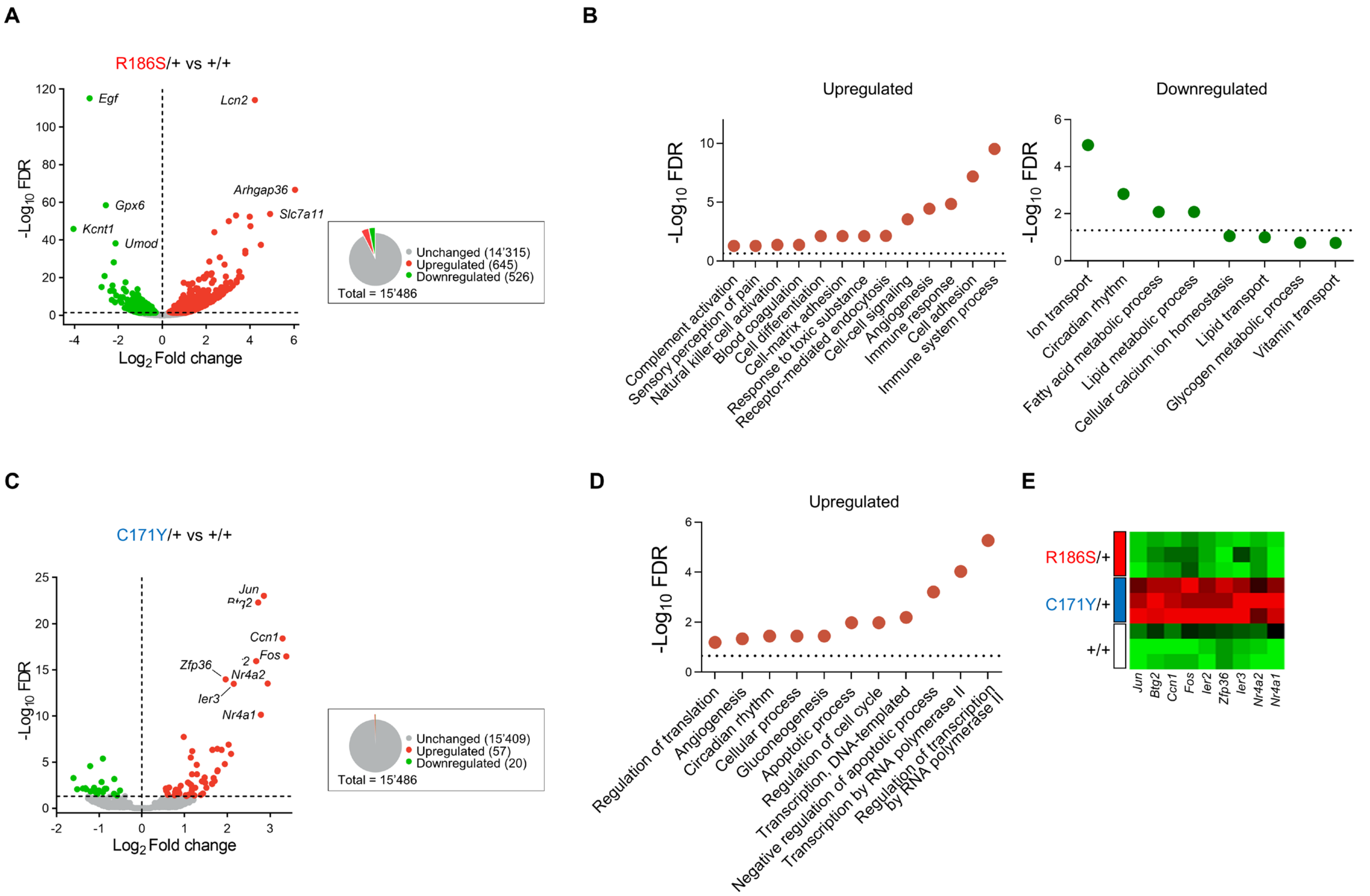

**Supplementary Figure 7: Differential expression and affected pathways in 4-month-old *Umod* KI kidneys.**

(**A**) Volcano plot showing differently expressed genes (DEGs) between +/+ and R186S/+ kidneys at 4 months. (**B**) Over-representation analysis (ORA) showing up- and downregulated biological processes in 4-months-old R186S/+ kidneys. (**C**) Volcano plot showing DEGs between C171Y/+ and +/+ kidneys at 4 months. Genes not significantly changed (FDR > 0.05) are shown in grey, whereas genes that are up- or downregulated in *Umod* KI are shown in red and green respectively. The total number of unchanged, up- and downregulated genes are summarized in the pie chart. (**D**) ORA showing upregulated biological processes in 4-month-old C171Y/+ kidneys. No significantly downregulated pathways were identified. (**e**) Heatmap of transcriptional regulation genes upregulated in 4-month-old C171Y/+ kidneys.

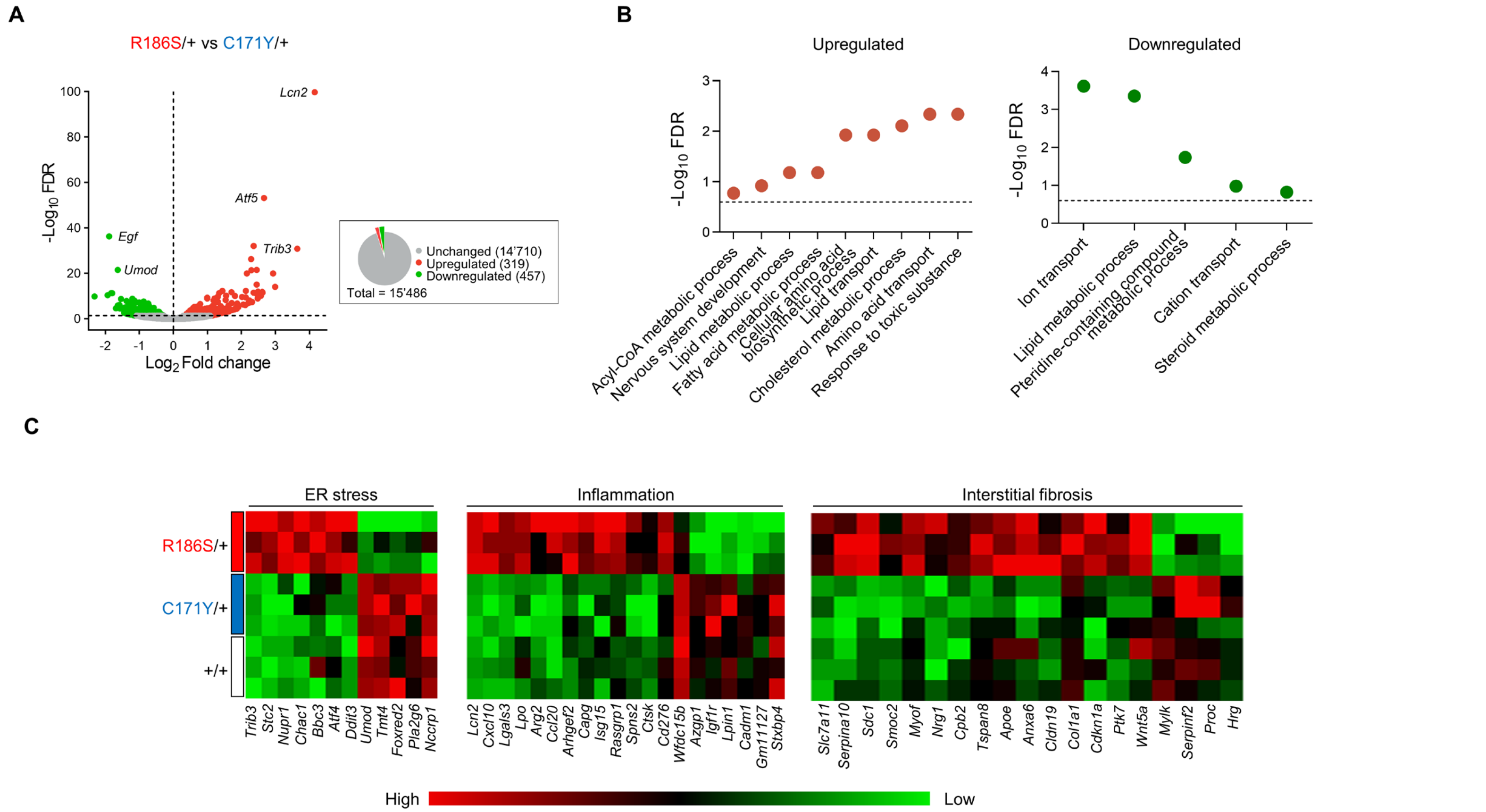

**Supplementary Figure 8: Disease progression signature in *Umod* KI kidneys.**

(**A**) Volcano plot showing differently expressed genes (DEGs) between R186S/+ and C171Y/+ kidneys at 1 month. Genes not significantly changed (FDR > 0.05) are shown in grey, whereas genes that are up- or downregulated in R186S/+ are shown in red and green respectively. The total number of unchanged, up- and downregulated genes are summarized in the pie chart. (**B**) Over-representation analysis (ORA) showing up- (red) and downregulated (green) biological processes in 1-month-old R186S/+ kidneys compared to C171Y/+. (**C**) Heat map of selected pathways involved in the disease progression of *Umod* KI mice at 1 month. R186S/+ kidneys showed upregulation of ER stress (*Atf4*, *Ddit3*, *Nupr1*, *Trib3*), increased expression of genes associated with inflammation (*Lcn2*, *Lgals3*) and fibrosis (*Serpina10*, *Col1a1*), whereas C171Y/+ kidneys were virtually indistinguishable from +/+.

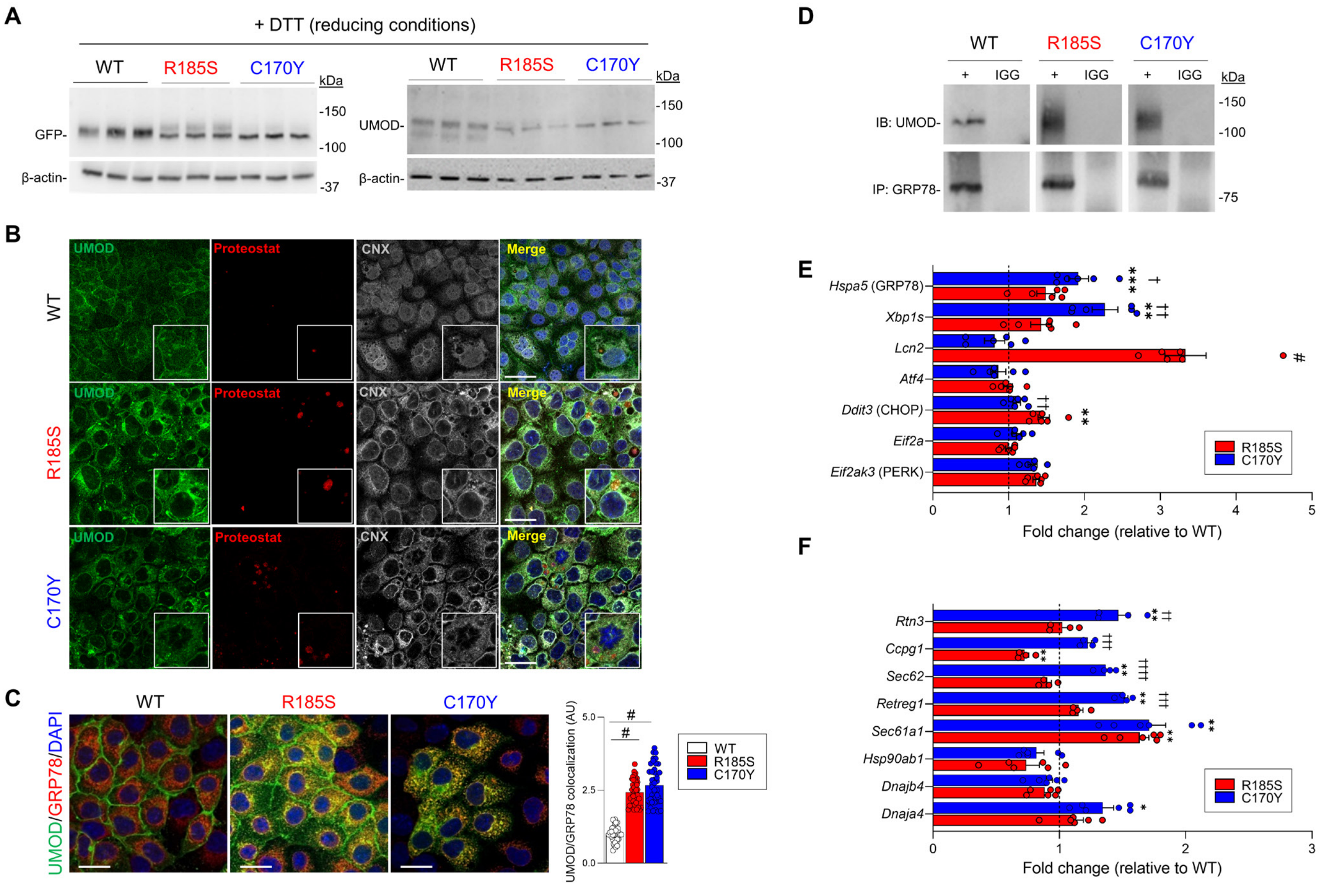

**Supplementary Figure 9: Distinct UMOD mutations trigger differential ER quality control responses.**
(**A**) Immunoblot analysis of GFP and UMOD in lysates from *UMOD-GFP* cells. Samples were run in reducing conditions. β-actin used as a loading control. (**B**) Immunofluorescence analysis of UMOD (green), Proteostat (red) and CNX (gray) in *UMOD-GFP* cells. Scale bar: 30 µm. (**C**) Immunofluorescence analysis of UMOD (green) and GRP78/BiP (red) in *UMOD-GFP* cells. Co-localization is expressed as arbitrary units (AU). Scale bar: 30 µm (n=33 to 44 cells per group). (**D**) Co-immunoprecipitation experiments in *UMOD-GFP* cells showing interaction between UMOD and GRP78/BiP. (**E**) RT-qPCR analysis of unfolded protein response (UPR) effectors in in *UMOD-GFP* cells. Values are expressed as relative to WT (black dotted line, n = 6 biological replicates). (**F**) RT-qPCR analysis of protein folding/degradation genes in *UMOD-GFP* cells. Values are expressed as relative to WT (black dotted line) (n =4 to 6 biological replicates). Bars indicate mean ± SEM. One-way ANOVA followed by Tukey's post-hoc test; *P < 0.05, **P < 0.01, #P < 0.0001 compared to WT; ††P < 0.01, †††P < 0.001, †††† P < 0.0001 R185S vs C170Y.

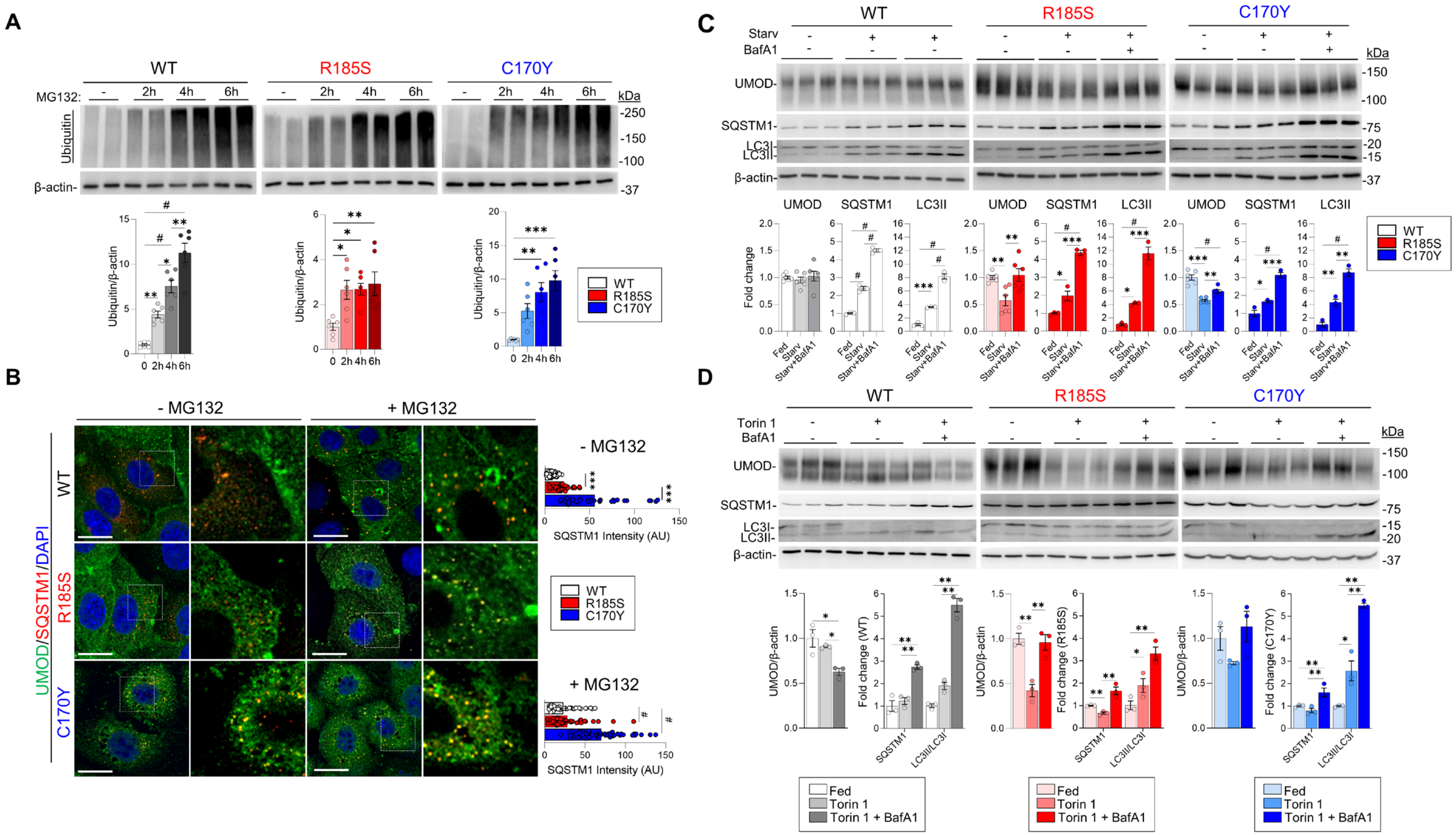

**Supplementary Figure 10: Autophagy induction enhances mutant UMOD clearance.**
(**A**) Immunoblot analysis of ubiquitin in *UMOD-GFP* cell lysates following MG123 time course. β-actin used as a loading control. Densitometry analysis relative to untreated cells (n = 6 biological replicates). (**B**) Representative immunofluorescence of UMOD (green) and SQSTM1 (red) in *UMOD-GFP* cells. Nuclei are counterstained with DAPI (blue). Scale bar: 15 μm (n = 30 cells per group). (**C**) Immunoblot analysis of UMOD, SQSTM1 and LC3 in *UMOD-GFP* cell lysates following starvation and Bafilomycin A1 treatment. β-actin used as a loading control. Densitometry analysis relative to fed cells (n=3 biological replicates). (**D**) Immunoblot analysis of UMOD, SQSTM1 and LC3 in *UMOD-GFP* cell lysates following Torin 1 and Bafilomycin A1 treatment. β-actin used as a loading control. Densitometry analysis relative to fed cells (n=3 biological replicates). Bars indicate mean ± SEM. One-way ANOVA followed by Tukey's post-hoc test; *$P < 0.05$, **$P < 0.01$, ***$P < 0.001$, #$P < 0.0001$.

**Supplementary Table 1: Clinical characteristics of p. (Arg185Ser) ADTKD-*UMOD* family.**

| | Gender | CKD (age) | Kidney failure (age) | Hyperuricemia (age) | Gout (age) |
|---|---|---|---|---|---|
| p.R185S III.1 | F | Y | Y (NA) | Y | Y (40y) |
| p.R185S III.2 | F | Y | Y (50y) | NA | NA |
| p.R185S IV.1 | M | Y | Y (43y) | Y | Y |
| p.R185S IV.2 | M | Y (32y) | Y (36y) | Y | Y (32y) |
| p.R185S IV.3 | M | Y (19y) | Y (28y) | Y | Y (9y) |
| p.R185S IV.4 | F | Y | N (41y) | Y (34y) | N |
| p.R185S IV.5 | F | Y | Y (38y) | Y | Y (18y) |
| p.R185S IV.6 | M | Y (38y) | Y (42y) | Y | Y (26y) |
| p.R185S V.1 | M | Y | N (24y) | NA | NA |
| p.R185S V.2 | M | Y | N (34y) | Y | Y (25y) |

M, male; F, female; Y, yes; N, no; NA, not available; CKD, chronic kidney disease; eGFR, estimated glomerular filtration rate.

**Supplementary Table 2: Clinical characteristics of p. (Cys170Tyr) ADTKD-*UMOD* families.**

| F1 | Gender | CKD (age) | Kidney failure (age) | Hyperuricemia | Gout (age) |
|---|---|---|---|---|---|
| p.C170Y I.1 | M | Y | Y (63y) | Y | Y |
| p.C170Y II.1 | M | Y (41y) | Y (58y; post-nephrectomy)* | Y | Y (55y) |
| p.C170Y III.1 | F | Y (43y) | N | Y | N |
| p.C170Y III.2 | F | Y | N (28y) | NA | NA |
| p.C170Y IV.1 | F | N (12y) | N | N | N |

| F2 | Gender | CKD (age) | Kidney failure (age) | Hyperuricemia | Gout (age) |
|---|---|---|---|---|---|
| p.C170Y I.1 | M | Y | Y (69y) | Y | Y |
| p.C170Y II.1 | F | Y | N (73y: eGFR 33mL/min; 86y: no ESKD) | Y | N |
| p.C170Y II.2 | F | Y | Y (84y) | Y | N |
| p.C170Y III.1 | F | Y (33y) | N (54y) | Y | N |

M, male; F, female; Y, yes; N, no; NA, not available; CKD, chronic kidney disease; eGFR, estimated glomerular filtration rate.
* Nephrectomy for clear cell carcinoma – abundant hematuria

**Supplementary Table 3: In silico analysis of selected *UMOD* missense variants.**

| Genomic coordinates (GRCh38) | Nucleotide change | Predicted amino acid change | Control & patient databases* | Pathogenicity predictions¥ | Comment | ACMG classification (PMID 25741868) |
|---|---|---|---|---|---|---|
| 16:20348792:C:T | c.509G>A | p.(Cys170Tyr) | 0¶ | Pathogenic computational verdict (7 path. vs. 4 ben. predictions), REVEL metascore: 0.63† | Not in ClinVar; located between EGF-like 3 domain and D8C UniProt classifies this variant as Pathogenic, associated with Tubulointerstitial kidney disease, autosomal dominant, 1. | **Likely Pathogenic (PM2, PP2, PP3, PP5, PP1)** |
| 16:20348748:G:T | c.553C>A | p.(Arg185Ser) | 0 | Pathogenic computational verdict (9 path. vs. 2 ben. predictions), REVEL metascore: 0.84† | Not in ClinVar; located between EGF-like 3 domain and D8C; inside a mutational hot-spot# UniProt classifies this variant as Pathogenic, associated with Tubulointerstitial kidney disease, autosomal dominant, 1. | **Pathogenic (PM1, PM5, PM2, PP2, PP3, PP5, PP1)** |

UMOD transcript: NM_001008389.3

* Includes the Genome Aggregation Database (gnomAD) (1), the UK Biobank (2), the Genomics England 100,000 Genomes Project (3)and the UK Rare Disease Registry (RaDaR) (https://ukkidney.org/rare-renal/radar).

¶ Has been reported before in 1 additional French family with slowly progressive CKD (2 individuals with 72y & 73y not yet in ESKD) (4)

¥ Generated using Varsome (5).

† REVEL, rare exome variant ensemble learner (6) (a score > 0.75 corresponds to a sensitivity of ~0.5 and a specificity of ~0.95 for pathogenic variants in the training dataset).

#Hot-spot of length 17 amino-acids has 9 missense/in-frame/non-synonymous variants (5 pathogenic, 3 uncertain, and 1 benign), which qualifies as a dense hot-spot (6).

Abbreviations: path., pathogenic; ben., benign; EGF-like, epidermal growth factor-like domain; D8C, cysteine-rich domain of unknown function.

**Supplementary Table 4: Clinical and biochemical parameters of *Umod*$^{C171Y}$ mice.**

| | 1 month | | | 4 months | | |
|---|---|---|---|---|---|---|
| **Parameter** | *Umod*$^{+/+}$ n=9 | *Umod*$^{C17Y/+}$ n=16 | *Umod*$^{C171Y/C171Y}$ n=5 | *Umod*$^{+/+}$ n=5 | *Umod*$^{C17Y/+}$ n=11 | *Umod*$^{C171Y/C171Y}$ n=7 |
| Body weight (g) | 19.7 ± 0.7 | 20.2 ± 0.6 | 19.1 ± 1.5 | 25.6 ± 1.5 | 28.3 ± 1.0 | 27.4 ± 1.5 |
| Water intake (µl/min/g body weight) | 0.23 ± 0.03 | 0.27 ± 0.01 | 0.29 ± 0.02 | 0.17 ± 0.02 | 0.14 ± 0.01 | 0.09 ± 0.02* |
| **Urine** | | | | | | |
| Diuresis (µl/min/g body weight) | 0.07 ± 0.02 | 0.05 ± 0.01 | 0.04 ± 0.01 | 0.05 ± 0.01 | 0.05 ± 0.01 | 0.05 ± 0.01 |
| $Na^+$ (g/g creat) | 5.6 ± 0.3 | 5.7 ± 0.2 | 7.2 ± 0.7 | 4.5 ± 0.6 | 6.6 ± 0.4* | 4.9 ± 0.7 |
| $K^+$ (g/g creat) | 19.5 ± 1.0 | 20.1 ± 0.6 | 23.6 ± 2.0 | 15.7 ± 0.9 | 19.2 ± 1.0 | 15.7 ± 1.0 |
| $Cl^-$ (g/g creat) | 14.4 ± 0.8 | 14.3 ± 0.4 | 17.9 ± 1.0* | 9.8 ± 0.8 | 12.6 ± 1.0 | 10.1 ± 0.8 |
| $Ca^{2+}$ (g/g creat) | 0.14 ± 0.02 | 0.14 ± 0.01 | 0.17 ± 0.02 | 0.17 ± 0.04 | 0.09 ± 0.01* | 0.17 ± 0.02 |
| $Mg^{2+}$ (g/g creat) | 1.32 ± 0.06 | 1.24 ± 0.08 | 1.39 ± 0.04 | 0.95 ± 0.04 | 1.03 ± 0.03 | 1.02 ± 0.07 |
| Creatinine (mg/dL) | 44 ± 3 | 43 ± 2 | 32 ± 3* | 41 ± 5 | 32 ± 2 | 35 ± 5 |
| $FE_{UA}$ (%) | 1.16 ± 0.26 | 0.97 ± 0.12 | 1.41 ± 0.34 | 0.59 ± 0.11 | - | - |
| Osmolality (mOsm/kg $H_2O$) | 1854 ± 85 | 1816 ± 93 | 1648 ± 132 | 1476 ± 184 | 1364 ± 109 | 1284 ± 188 |
| **Plasma** | n=9 | n=21 | n=9 | n=16 | n=20 | n=24 |
| $Na^+$ (mmol/l) | 147 ± 2 | 146 ± 1 | 148 ± 1 | 147 ± 1 | 149 ± 1 | 149 ± 1 |
| $Cl^-$ (mmol/l) | 108 ± 1 | 108 ± 2 | 108 ± 2 | 111 ± 1 | 110 ± 0.8 | 111 ± 1 |
| $Ca^{2+}$ (mmol/l) | 2.8 ± 0.04 | 2.8 ± 0.05 | 2.8 ± 0.6 | 2.5 ± 0.05 | 2.5 ± 0.03 | 2.50 ± 0.3 |
| Creatinine (mg/dl) | 0.14 ± 0.02 | 0.13 ± 0.01 | 0.15 ± 0.03 | 0.12 ± 0.01 | 0.13 ± 0.01 | 0.15 ± 0.01 |
| BUN (mg/dl) | 21 ± 2 | 19 ± 1 | 21 ± 1 | 20 ± 2 | 25 ± 1* | 29 ± 1*** |
| Uric acid (mg/dl) | 6.4 ± 0.4 | 5.9 ± 0.3 | 5.2 ± 0.4 | 5.2 ± 0.5 | 4.6 ± 0.4 | 4.0 ± 0.3 |
| Osmolality (mOsm/kg $H_2O$) | 351 ± 3 | 343 ± 3 | 347 ± 4 | 335 ± 3 | 342 ± 2* | 343 ± 2 |

Values are presented as average ± SEM. *$P < 0.05$, **$P < 0.01$, ***$P < 0.001$, #$P < 0.0001$ versus age matched Umod$^{+/+}$ mice. n: number of animals, FEUA: fractional excretion of uric acid, BUN: blood urea nitrogen.

**Supplementary Table 5: Clinical and biochemical parameters of *Umod*$^{R186S}$ mice.**

| | 1 month | | | 4 months | | |
|---|---|---|---|---|---|---|
| **Parameter** | *Umod*$^{+/+}$ n=15 | *Umod*$^{R186S/+}$ n=18 | *Umod*$^{R186S/R186S}$ n=12 | *Umod*$^{+/+}$ n=10 | *Umod*$^{R186S/+}$ n=13 | *Umod*$^{R186S/R186S}$ n=7 |
| Body weight (g) | 16.5 ± 0.6 | 16.4 ± 0.6 | 16.4 ± 0.6 | 26.3 ± 1.4 | 23.3 ± 1.0 | 26.5 ± 0.9 |
| Water intake (µl/min/g body weight) | 0.29 ± 0.02 | 0.27 ± 0.02 | 0.28 ± 0.03 | 0.15 ± 0.01 | 0.31 ± 0.02*** | 0.44 ± 0.04*** |
| **Urine** | | | | | | |
| Diuresis (µl/min/g BW) | 0.041 ± 0.006 | 0.059 ± 0.009 | 0.14 ± 0.04* | 0.046 ± 0.007 | 0.12 ± 0.01*** | 0.23 ± 0.02*** |
| $Na^+$ (g/g creat) | 5.8 ± 0.6 | 6.3 ± 0.7 | 6.4 ± 0.8 | 5.6 ± 0.3 | 5.7 ± 0.5 | 4.7 ± 0.6 |
| $K^+$ (g/g creat) | 27.7 ± 1.3 | 23.8 ± 0.9* | 27.06 ± 1.1 | 19.7 ± 1.2 | 21.4 ± 1.16 | 19.4 ± 0.6 |
| $Cl^-$ (g/g creat) | 19.8 ± 1.2 | 17.5 ± 1.0 | 21.3 ± 1.1 | 14.5 ± 1.1 | 15.9 ± 1.0 | 14.2 ± 0.7 |
| $Ca^{2+}$ (g/g creat) | 0.15 ± 0.02 | 0.28 ± 0.04* | 0.24 ± 0.03* | 0.08 ± 0.07 | 0.21 ± 0.02# | 0.20 ± 0.03*** |
| $Mg^{2+}$ (g/g creat) | 2.2 ± 0.9 | 2.0 ± 0.1 | 2.3 ± 0.1 | 1.3 ± 0.1 | 1.41 ± 0.13 | 1.65 ± 0.08 |
| Creatinine (mg/dL) | 43 ± 3 | 34 ± 3 | 29 ± 3** | 50 ± 2 | 25 ± 2# | 15 ± 1# |
| $FE_{UA}$ (%) | 1.16 ± 0.26 | 1.18 ± 0.31 | 0.72 ± 0.36 | 0.25 ± 0.1 | 0.74 ± 0.4*** | 0.16 ± 0.05** |
| Osmolality (mOsm/kg $H_2O$) | 2227 ± 186 | 1647 ± 168* | 1440 ± 152** | 2012 ± 132 | 958 ± 58# | 583 ± 20# |
| **Plasma** | n=9 | n=6 | n=13 | n=23 | n=24 | n=14 |
| $Na^+$ (mmol/l) | 149 ± 2 | 149 ± 1 | 149 ± 1 | 147 ± 1 | 149 ± 1* | 153 ± 1*** |
| $Cl^-$ (mmol/l) | 109 ± 1 | 109 ± 1 | 108 ± 1 | 119 ± 1 | 108 ± 1 | 107 ± 1* |
| $Ca^{2+}$ (mmol/l) | 2.72 ± 0.05 | 2.63 ± 0.06 | 2.64 ± 0.06 | 2.50 ± 0.03 | 2.51 ± 0.03 | 2.65 ± 0.04* |
| Creatinine (mg/dl) | 0.17 ± 0.04 | 0.12 ± 0,01 | 0.15 ± 0.01 | 0.11 ± 0.01 | 0.14 ± 0.01* | 0.18 ± 0.01*** |
| BUN (mg/dl) | 19 ± 1 | 33 ± 4*** | 43 ± 4# | 21 ± 1 | 55 ± 2# | 70 ± 2# |
| Uric acid (mg/dl) | 6.6 ± 0.7 | 5.2 ± 1.1 | 5.9 ± 0.5 | 5.4 ± 0.3 | 4.8 ± 0.4 | 5.1 ± 0.8 |
| Osmolality (mOsm/kg $H_2O$) | 353 ± 4 | 351 ± 4 | 356 ± 3 | 340 ± 2 | 355 ± 2# | 366 ± 3# |

Values are presented as average ±SEM. *$P < 0.05$, **$P < 0.01$, ***$P < 0.001$ versus age matched *Umod*$^{+/+}$ mice. $FE_{UA}$, Fractional excretion of uric acid; n, Number of animals; BUN, Blood urea nitrogen.

**Supplementary Table 6: Clinical and biochemical parameters of *Umod*$^{R186S/-}$ mice.**

| Parameter | 1 month old | | | 4 months old | | |
|---|---|---|---|---|---|---|
| | *Umod*$^{+/+}$ n=11 | *Umod*$^{R186S/+}$ n=10 | *Umod*$^{R186S/-}$ n=9 | *Umod*$^{+/+}$ n=8 | *Umod*$^{R186S/+}$ n=7 | *Umod*$^{R186S/-}$ n=13 |
| Body weight (g) | 16.1 ± 0.7 | 15.6 ± 0.6 | 16.6 ± 0.6 | 29.2 ± 1.7 | 26.6 ± 1.3 | 29.4 ± 0.6 |
| Water intake (µl/min/g body weight) | 0.36 ± 0.03 | 0.30 ± 0.01 | 0.33 ± 0.02 | 0.15 ± 0.02 | 0.29 ± 0.02*** | 0.25 ± 0.02** |
| **Urine** | | | | | | |
| Diuresis (µl/min/g BW) | 0.048 ± 0.008 | 0.057 ± 0.009 | 0.04 ± 0.01 | 0.03 ± 0.01 | 0.13 ± 0.02*** | 0.11 ± 0.01# |
| $Na^+$ (g/g creat) | 8.3 ± 0.5 | 9.0 ± 0.4 | 9.0 ± 0.6 | 3.2 ± 0.5 | 4.0 ± 0.3 | 3.9 ± 0.3 |
| $K^+$ (g/g creat) | 27.8 ± 0.9 | 30.0 ± 0.8 | 30.8 ± 1.4 | 18.7 ± 1.2 | 19.1 ± 1.2 | 18.5 ± 0.8 |
| $Cl^-$ (g/g creat) | 23.9 ± 0.8 | 25.5 ± 1.0 | 26.3 ± 1.4 | 10.8 ± 0.4 | 12.4 ± 1.0 | 11.6 ± 0.8 |
| $Ca^{2+}$ (g/g creat) | 0.23 ± 0.03 | 0.27 ± 0.02 | 0.28 ± 0.03 | 0.14 ± 0.02 | 0.22 ± 0.01** | 0.22 ± 0.01** |
| $Mg^{2+}$ (g/g creat) | 2.0 ± 0.2 | 2.2 ± 0.1 | 2.2 ± 0.2 | 1.2 ± 0.1 | 1.43 ± 0.07 | 1.32 ± 0.09 |
| Creatinine (mg/dL) | 35.2 ± 3.0 | 30.7 ± 1.4 | 37.2 ± 4.1 | 44.2 ± 5.5 | 21.7 ± 2.3** | 20.3 ± 1.1# |
| $FE_{UA}$ (%) | - | - | - | 0.55 ± 0.09 | 0.14 ± 0.02** | 0.16 ± 0.03** |
| Osmolality (mOsm/kg $H_2O$) | 1855 ± 120 | 1766 ± 81 | 2091 ± 217 | 1577 ± 201 | 771 ± 42** | 731 ± 29# |
| **Plasma** | n=13 | n=10 | n=11 | n=14 | n=15 | n=20 |
| $Na^+$ (mmol/l) | 145.9 ± 0.8 | 145.3 ± 0.8 | 145.1 ± 0.7 | 147.3 ± 0.7 | 149.1 ± 0.5* | 148.2 ± 0.5 |
| $Cl^-$ (mmol/l) | 107.8 ± 0,7 | 107.1 ± 0.5 | 110.3 ± 0.9 | 110.0 ± 1.1 | 107.1 ± 0.7* | 106 .6 ± 0.3** |
| $Ca^{2+}$ (mmol/l) | 2.67 ± 0.07 | 2.82 ± 0.05 | 2.77 ± 0.05 | 2.46 ± 0.05 | 2.62 ± 0.03** | 2.62 ± 0.03** |
| Creatinine (mg/dl) | 0.15 ± 0.02 | 0.142 ± 0.02 | 0.17 ± 0.03 | 0.10 ± 0.02 | 0.17 ± 0.02* | 0.16 ± 0.01* |
| BUN (mg/dl) | 19 ± 1 | 32 ± 2# | 32 ± 1.# | 20 ± 1.0 | 53 ± 4 # | 56 ± 1.# |
| Uric acid (mg/dl) | 3.3 ± 0.5 | 4.0 ± 0.9 | 4.8 ± 0.5 | 4.1 ± 0.3 | 4.1 ± 0.4 | 3.8 ± 0.4 |
| Osmolality (mOsm/kg $H_2O$) | 341 ± 4 | 353 ± 3 | 355 ± 4* | 338 ± 3 | 353 ± 3** | 357 ± 3# |

Values are presented as average ±SEM. *$P < 0.05$, **$P < 0.01$, ***$P < 0.001$, #$P < 0.0001$ versus age-matched *Umod*$^{+/+}$ mice or Ϯ $P \leq 0.05$ versus age-matched *Umod*$^{R186S/+}$ mice. n: number of animals, BUN: Blood Urea Nitrogen.

### Supplementary Table 7: Top 50 DEGs in *Umod*$^{R186S/+}$ kidneys at 1 month.

| Symbol | Gene name | Fold change (log2) | FDR (-log10) |
|---|---|---|---|
| *Lcn2* | Lipocalin 2 | 4.09 | 100.44 |
| *Atf5* | Activating transcription factor 5 | 2.61 | 52.02 |
| *Trib3* | Tribbles pseudokinase 3 | 4.04 | 37.43 |
| *Asns* | Asparagine synthetase | 2.39 | 33.55 |
| *Mthfd2* | Methylenetetrahydrofolate dehydrogenase (NAD+ dependent), methenyltetrahydrofolate cyclohydrolase | 2.31 | 27.30 |
| *Akr1b8* | Aldo-keto reductase family 1, member B8 | 2.21 | 20.17 |
| *Cxcl10* | Chemokine (C-X-C motif) ligand 10 | 2.92 | 19.70 |
| *Stc2* | Stanniocalcin 2 | 2.29 | 19.52 |
| *Aldh18a1* | Aldehyde dehydrogenase 18 family, member A1 | 2.08 | 18.88 |
| *Angptl6* | Angiopoietin-like 6 | 2.33 | 14.69 |
| *Aldh1l2* | Aldehyde dehydrogenase 1 family, member L2 | 1.87 | 14.36 |
| *Slc7a11* | Solute carrier family 7 (cationic amino acid transporter, y+ system), member 11 | 2.56 | 13.53 |
| *Mt2* | Metallothionein 2 | 1.76 | 13.17 |
| *Slc7a3* | Solute carrier family 7 (cationic amino acid transporter, y+ system), member 3 | 2.62 | 12.81 |
| *Nupr1* | Nuclear protein transcription regulator 1 | 1.45 | 12.61 |
| *Cbr3* | Carbonyl reductase 3 | 2.15 | 12.12 |
| *Arhgap36* | Rho GTPase activating protein 36 | 2.76 | 11.99 |
| *Soat2* | Sterol O-acyltransferase 2 | 2.56 | 11.48 |
| *Pappa* | Pregnancy-associated plasma protein A | 1.91 | 11.33 |
| *Slc38a1* | Solute carrier family 38, member 1 | 1.40 | 11.25 |
| *Wnt10a* | Wingless-type MMTV integration site family, member 10A | 2.65 | 10.75 |
| *Gabrp* | Gamma-aminobutyric acid (GABA) A receptor, pi | 2.51 | 10.74 |
| *Loxl4* | Lysyl oxidase-like 4 | 1.66 | 9.76 |
| *Lgals3* | Lectin, galactose binding, soluble 3 | 1.24 | 9.48 |
| *Fam129a* | Family with sequence similarity 129, member A | 1.32 | 8.13 |
| *Egf* | Epidermal growth factor | -2.00 | 40.86 |
| *Umod* | Uromodulin | -1.74 | 24.75 |
| *Car3* | Carbonic anhydrase 3 | -2.01 | 14.36 |
| *Gm36797* | Predicted gene, 36797 | -2.65 | 13.46 |
| *Gm32960* | Predicted gene, 32960 | -1.74 | 13.17 |
| *Kcnt1* | Potassium channel, subfamily T, member 1 | -1.87 | 9.55 |
| *Wfdc15b* | WAP four-disulfide core domain 15B | -1.23 | 7.59 |
| *Azgp1* | Aalpha-2-glycoprotein 1, zinc | -1.47 | 6.76 |
| *Dusp15* | Dual specificity phosphatase-like 15 | -1.46 | 6.47 |
| *Cyp2a4* | Cytochrome P450, family 2, subfamily a, polypeptide 4 | -1.04 | 6.32 |
| *Slc6a6* | Solute carrier family 6 (neurotransmitter transporter, taurine), member 6 | -0.63 | 5.88 |
| *Atp8b4* | ATPase, class I, type 8B, member 4 | -1.56 | 5.67 |
| *Fam107a* | Family with sequence similarity 107, member A | -0.99 | 5.53 |
| *Slco1a6* | Solute carrier organic anion transporter family, member 1a6 | -0.74 | 4.95 |
| *Tmem207* | Transmembrane protein 207 | -1.39 | 4.87 |
| *BC040756* | cDNA sequence BC040756 | -0.87 | 4.74 |
| *Pex5l* | Peroxisomal biogenesis factor 5-like | -1.48 | 4.49 |
| *Esrrb* | Estrogen related receptor, beta | -0.97 | 4.49 |
| *Cwh43* | Cell wall biogenesis 43 C-terminal homolog | -0.78 | 4.49 |
| *6330410L21Rik* | RIKEN cDNA 6330410L21 gene | -1.35 | 4.41 |
| *Ugt8a* | UDP galactosyltransferase 8A | -0.85 | 4.41 |
| *Omd* | Osteomodulin | -1.12 | 4.19 |
| *Rnf169* | Ring finger protein 169 | -0.88 | 4.18 |
| *Dok2* | Docking protein 2 | -1.13 | 4.13 |
| *Perm1* | PPARGC1 and ESRR induced regulator, muscle 1 | -1.36 | 4.09 |

FDR: False discovery rate-adjusted *P* value (Benjamini-Hochberg correction).

**Supplementary Table 8: Top 50 DEGs in *Umod*$^{R186S/+}$ kidneys at 4 months.**

| Symbol | Gene name | Fold change (log2) | FDR (-log10) |
|---|---|---|---|
| *Lcn2* | Lipocalin 2 | 4.23 | 114.29 |
| *Arhgap36* | Rho GTPase activating protein 36 | 6.04 | 66.64 |
| *Slc7a11* | Solute carrier family 7 (cationic amino acid transporter, y+ system), member 11 | 4.91 | 53.68 |
| *Akr1b8* | Aldo-keto reductase family 1, member B8 | 3.36 | 52.89 |
| *Nefl* | Neurofilament, light polypeptide | 3.98 | 52.36 |
| *Dpt* | Dermatopontin | 3.03 | 49.97 |
| *Cbr3* | Carbonyl reductase 3 | 4.02 | 47.22 |
| *Atf5* | Activating transcription factor 5 | 2.38 | 44.14 |
| *Ppp2r2c* | Protein phosphatase 2, regulatory subunit B, gamma | 4.49 | 37.30 |
| *Trib3* | Tribbles pseudokinase 3 | 3.78 | 34.16 |
| *B4galnt2* | Beta-1,4-N-acetyl-galactosaminyl transferase 2 | 3.78 | 32.79 |
| *Aldh18a1* | Aldehyde dehydrogenase 18 family, member A1 | 2.62 | 30.76 |
| *Asns* | Asparagine synthetase | 2.14 | 28.48 |
| *Stc2* | Stanniocalcin 2 | 2.84 | 26.86 |
| *Gabrp* | Gamma-aminobutyric acid (GABA) A receptor, pi | 3.51 | 23.20 |
| *Aldh1l2* | Aldehyde dehydrogenase 1 family, member L2 | 2.25 | 22.19 |
| *Abcc3* | ATP-binding cassette, sub-family C (CFTR/MRP), member 3 | 2.21 | 21.80 |
| *Smoc2* | SPARC related modular calcium binding 2 | 2.04 | 21.77 |
| *Astn2* | Astrotactin 2 | 2.22 | 21.40 |
| *Fcrls* | Fc receptor-like S, scavenger receptor | 3.50 | 20.52 |
| *Kif1a* | Kinesin family member 1A | 2.90 | 20.50 |
| *Lyz2* | Lysozyme 2 | 1.98 | 20.30 |
| *Gm32857* | Predicted gene, 32857 | 3.62 | 20.28 |
| *Mrc1* | Mannose receptor, C type 1 | 1.89 | 20.28 |
| *Mthfd2* | Methylenetetrahydrofolate dehydrogenase (NAD+ dependent), methenyltetrahydrofolate cyclohydrolase | 1.92 | 19.46 |
| *Egf* | Epidermal growth factor | -3.30 | 115.22 |
| *Gpx6* | Glutathione peroxidase 6 | -2.57 | 58.29 |
| *Kcnt1* | Potassium channel, subfamily T, member 1 | -4.04 | 45.73 |
| *Umod* | Uromodulin | -2.13 | 38.09 |
| *Wfdc15b* | WAP four-disulfide core domain 15B | -2.21 | 28.02 |
| *Tmem207* | Transmembrane protein 207 | -2.62 | 20.68 |
| *Aldoc* | Aldolase C, fructose-bisphosphate | -1.67 | 17.42 |
| *Dusp15* | Dual specificity phosphatase-like 15 | -2.27 | 15.75 |
| *Gm36797* | Predicted gene, 36797 | -2.76 | 14.99 |
| *Gm32960* | Predicted gene, 32960 | -1.76 | 14.00 |
| *Ppp1r1a* | Protein phosphatase 1, regulatory inhibitor subunit 1A | -1.13 | 13.72 |
| *Pex5l* | Peroxisomal biogenesis factor 5-like | -2.36 | 12.85 |
| *Perm1* | PPARGC1 and ESRR induced regulator, muscle 1 | -2.21 | 12.77 |
| *Lrrc66* | Leucine rich repeat containing 66 | -1.65 | 12.49 |
| *Ckb* | Creatine kinase, brain | -1.36 | 11.45 |
| *Ank2* | Ankyrin 2, brain | -1.18 | 10.87 |
| *Mfsd4a* | Major facilitator superfamily domain containing 4A | -1.10 | 10.75 |
| *6330410L21Rik* | RIKEN cDNA 6330410L21 gene | -2.09 | 10.48 |
| *Pcsk6* | Proprotein convertase subtilisin/kexin type 6 | -1.27 | 10.45 |
| *Slc6a12* | Solute carrier family 6 (neurotransmitter transporter, betaine/GABA), member 12 | -1.90 | 9.48 |
| *Spag5* | Sperm associated antigen 5 | -1.78 | 9.48 |
| *Clcnka* | Chloride channel, voltage-sensitive Ka | -1.29 | 9.46 |
| *Gcgr* | Glucagon receptor | -1.19 | 9.23 |
| *Pla1a* | Phospholipase A1 member A | -1.00 | 8.76 |
| *Aqp4* | Aquaporin 4 | -1.37 | 8.33 |

FDR: False discovery rate-adjusted *P* value (Benjamini-Hochberg correction).

### Supplementary Table 9: Top 50 DEGs in *Umod*$^{C171Y/+}$ kidneys at 4 months.

| Symbol | Gene name | Fold change (log2) | FDR (-log10) |
|---|---|---|---|
| *Jun* | Jun proto-oncogene | 2.86 | 23.00 |
| *Btg2* | BTG anti-proliferation factor 2 | 2.72 | 22.27 |
| *Ccn1* | Cellular communication network factor 1 | 3.29 | 18.39 |
| *Fos* | FBJ osteosarcoma oncogene | 3.38 | 16.46 |
| *Ier2* | Immediate early response 2 | 2.68 | 15.92 |
| *Zfp36* | Zinc finger protein 36 | 1.96 | 13.97 |
| *Nr4a2* | Nuclear receptor subfamily 4, group A, member 2 | 2.94 | 13.51 |
| *Ier3* | Immediate early response 3 | 2.15 | 13.47 |
| *Nr4a1* | Nuclear receptor subfamily 4, group A, member 1 | 2.79 | 10.13 |
| *Fosb* | FBJ osteosarcoma oncogene B | 2.03 | 6.90 |
| *Snord14e* | Small nucleolar RNA, C/D box 14° | 1.77 | 6.43 |
| *Csrnp1* | Cysteine-serine-rich nuclear protein 1 | 1.65 | 6.32 |
| *Gdf15* | Growth differentiation factor 15 | 1.87 | 6.32 |
| *Ccn2* | Cellular communication network factor 2 | 1.18 | 6.18 |
| *Gm30591* | Predicted gene, 30591 | 2.09 | 5.88 |
| *Tob1* | Transducer of ErbB-2.1 | 1.14 | 5.47 |
| *Egr1* | Early growth response 1 | 1.94 | 4.77 |
| *Gm17971* | Predicted gene, 17971 | 1.28 | 4.69 |
| *Snord14d* | Small nucleolar RNA, C/D box 14D | 1.77 | 4.11 |
| *Egr3* | Early growth response 3 | 1.76 | 3.99 |
| *Rasd1* | RAS, dexamethasone-induced 1 | 1.29 | 3.68 |
| *Dusp6* | Dual specificity phosphatase 6 | 1.17 | 3.67 |
| *Ighv5-9* | Immunoglobulin heavy variable 5-9 | 1.47 | 3.31 |
| *Ch25h* | Cholesterol 25-hydroxylase | 1.68 | 3.27 |
| *Ighg2b* | Immunoglobulin heavy constant gamma 2B | 1.70 | 2.93 |
| *mt-Tl1* | tRNA leucine 1, mitochondrial | -1.21 | 4.57 |
| *Gm30238* | predicted gene, 30238 | -1.60 | 3.27 |
| *Igkv13-85* | immunoglobulin kappa chain variable 13-85 | -1.12 | 2.21 |
| *Cyp2d26* | cytochrome P450, family 2, subfamily d, polypeptide 26 | -1.32 | 2.16 |
| *Lpl* | lipoprotein lipase | -1.38 | 2.14 |
| *Synpr* | synaptoporin | -1.30 | 2.06 |
| *Mir6236* | microRNA 6236 | -1.51 | 2.05 |
| *1700067K01Rik* | RIKEN cDNA 1700067K01 gene | -1.01 | 1.91 |
| *Atp1a2* | ATPase, Na+/K+ transporting, alpha 2 polypeptide | -1.20 | 1.82 |
| *Itih1* | inter-alpha trypsin inhibitor, heavy chain 1 | -1.09 | 1.76 |
| *Apoc1* | apolipoprotein C-I | -1.04 | 1.55 |
| *Adig* | adipogenin | -0.91 | 5.38 |
| *Slc5a1* | solute carrier family 5 (sodium/glucose cotransporter), member 1 | -0.64 | 3.17 |
| *Rn18s-rs5* | 18s RNA, related sequence 5 | -0.94 | 2.83 |
| *LOC108167485* | putative uncharacterized protein FLJ37770 pseudogene | -0.87 | 2.11 |
| *Gm17597* | predicted gene, 17597 | -0.83 | 2.11 |
| *Mri1* | methylthioribose-1-phosphate isomerase 1 | -0.51 | 1.92 |
| *Gm34472* | predicted gene, 34472 | -0.91 | 1.73 |
| *Gm28023* | predicted gene, 28023 | -0.65 | 1.56 |
| *Tcf24* | transcription factor 24 | -0.58 | 1.37 |
| *Ugt3a1* | UDP glycosyltransferases 3 family, polypeptide A1 | -0.39 | 1.29 |
| *Cfd* | complement factor D (adipsin) | -1.26 | 1.25 |
| *AI429214* | expressed sequence AI429214 | -0.75 | 1.25 |
| *AA536875* | expressed sequence AA536875 | -0.92 | 1.15 |
| *Gm15564* | predicted gene 15564 | -1.22 | 1.15 |

FDR: False discovery rate-adjusted *P* value (Benjamini-Hochberg correction).

**Supplementary Table 10: Top 50 DEGs in *Umod*$^{R186S/+}$ compared to *Umod*$^{C171Y/+}$ kidneys at 1 month.**

| Symbol | Gene name | Fold change (log2) | FDR (-log10) |
|---|---|---|---|
| *Lcn2* | Lipocalin 2 | 4.16 | 99.72 |
| *Atf5* | Activating transcription factor 5 | 2.67 | 53.11 |
| *Asns* | Asparagine synthetase | 2.36 | 32.01 |
| *Trib3* | Tribbles pseudokinase 3 | 3.65 | 30.80 |
| *Mthfd2* | Methylenetetrahydrofolate dehydrogenase (NAD+ dependent), methenyltetrahydrofolate cyclohydrolase | 2.28 | 26.20 |
| *Stc2* | Stanniocalcin 2 | 2.45 | 21.46 |
| *Akr1b8* | Aldo-keto reductase family 1, member B8 | 2.29 | 21.33 |
| *Cxcl10* | Chemokine (C-X-C motif) ligand 10 | 2.94 | 19.82 |
| *Aldh18a1* | Aldehyde dehydrogenase 18 family, member A1 | 2.16 | 19.82 |
| *Cbr3* | Carbonyl reductase 3 | 2.45 | 14.95 |
| *Arhgap36* | Rho GTPase activating protein 36 | 2.99 | 13.98 |
| *Nupr1* | Nuclear protein transcription regulator 1 | 1.44 | 12.23 |
| *Angptl6* | Angiopoietin-like 6 | 2.13 | 12.16 |
| *Samd5* | Sterile alpha motif domain containing 5 | 2.49 | 11.82 |
| *Soat2* | Sterol O-acyltransferase 2 | 2.62 | 11.70 |
| *Aldh1l2* | Aldehyde dehydrogenase 1 family, member L2 | 1.71 | 11.70 |
| *Gabrp* | Gamma-aminobutyric acid (GABA) A receptor, pi | 2.62 | 11.40 |
| *Slc7a11* | Solute carrier family 7 (cationic amino acid transporter, y+ system), member 11 | 2.36 | 11.40 |
| *Lgals3* | Lectin, galactose binding, soluble 3 | 1.34 | 11.16 |
| *Slc7a3* | Solute carrier family 7 (cationic amino acid transporter, y+ system), member 3 | 2.44 | 10.98 |
| *Wnt10a* | Wingless-type MMTV integration site family, member 10A | 2.58 | 10.07 |
| *Mt2* | Metallothionein 2 | 1.55 | 10.03 |
| *Pappa* | Pregnancy-associated plasma protein A | 1.81 | 9.98 |
| *Slc38a1* | Solute carrier family 38, member 1 | 1.31 | 9.58 |
| *Serpina10* | Serine (or cysteine) peptidase inhibitor, clade A (alpha-1 antiproteinase, antitrypsin), member 10 | 2.02 | 9.52 |
| *Egf* | Epidermal growth factor | -1.89 | 36.25 |
| *Umod* | Uromodulin | -1.63 | 21.44 |
| *Azgp1* | Alpha-2-glycoprotein 1, zinc | -1.83 | 11.19 |
| *Car3* | Carbonic anhydrase 3 | -1.80 | 11.16 |
| *Kcnt1* | Potassium channel, subfamily T, member 1 | -1.93 | 10.25 |
| *Gm36797* | Predicted gene, 36797 | -2.32 | 9.74 |
| *Gm34472* | Predicted gene, 34472 | -1.21 | 8.77 |
| *Gm34861* | Predicted gene, 34861 | -1.54 | 8.63 |
| *Cyp2a4* | Cytochrome P450, family 2, subfamily a, polypeptide 4 | -1.14 | 7.88 |
| *Tc2n* | Tandem C2 domains, nuclear | -1.37 | 7.77 |
| *Agps* | Alkylglycerone phosphate synthase | -1.01 | 7.08 |
| *Ksr2* | Kinase suppressor of ras 2 | -1.10 | 7.05 |
| *Wfdc15b* | WAP four-disulfide core domain 15B | -1.18 | 7.01 |
| *Gm32960* | Predicted gene, 32960 | -1.30 | 6.72 |
| *Trpm6* | Transient receptor potential cation channel, subfamily M, member 6 | -1.26 | 6.64 |
| *Atp8b4* | ATPase, class I, type 8B, member 4 | -1.58 | 5.95 |
| *Syn3* | Synapsin III | -1.33 | 5.87 |
| *Col19a1* | Collagen, type XIX, alpha 1 | -1.60 | 5.79 |
| *Idi1* | Isopentenyl-diphosphate delta isomerase | -1.21 | 5.77 |
| *Slco1a1* | Solute carrier organic anion transporter family, member 1a1 | -1.63 | 5.67 |
| *Ubiad1* | UbiA prenyltransferase domain containing 1 | -1.26 | 5.31 |
| *Dusp15* | Dual specificity phosphatase-like 15 | -1.33 | 5.23 |
| *Gm38481* | Predicted gene, 38481 | -1.22 | 4.87 |
| *Gm7537* | Predicted gene 7537 | -1.27 | 4.81 |
| *Perm1* | PPARGC1 and ESRR induced regulator, muscle 1 | -1.43 | 4.80 |

FDR: False discovery rate-adjusted *P* value (Benjamini-Hochberg correction).

### Supplementary Table 11: Top 50 DEGs in *Umod*$^{R186S/+}$ compared to *Umod*$^{C171Y/+}$ at 4 months.

| Symbol | Gene name | Fold change (log2) | FDR (-log10) |
|---|---|---|---|
| *Lcn2* | Lipocalin 2 | 4.73 | 132.87 |
| *Akr1b8* | Aldo-keto reductase family 1, member B8 | 3.95 | 70.59 |
| *Arhgap36* | Rho GTPase activating protein 36 | 5.99 | 65.97 |
| *Slc7a11* | Solute carrier family 7 (cationic amino acid transporter, y+ system), member 11 | 4.76 | 51.56 |
| *Dpt* | Dermatopontin | 2.99 | 49.40 |
| *Nefl* | Neurofilament, light polypeptide | 3.41 | 42.31 |
| *Trib3* | Tribbles pseudokinase 3 | 4.00 | 37.82 |
| *Cbr3* | Carbonyl reductase 3 | 3.34 | 35.67 |
| *Atf5* | Activating transcription factor 5 | 2.10 | 35.22 |
| *Ppp2r2c* | Protein phosphatase 2, regulatory subunit B, gamma | 4.32 | 35.17 |
| *B4galnt2* | Beta-1,4-N-acetyl-galactosaminyl transferase 2 | 3.62 | 30.75 |
| *Asns* | Asparagine synthetase | 2.21 | 30.44 |
| *Gabrp* | Gamma-aminobutyric acid (GABA) A receptor, pi | 4.00 | 29.49 |
| *Aldh18a1* | Aldehyde dehydrogenase 18 family, member A1 | 2.32 | 25.09 |
| *Gsta4* | Glutathione S-transferase, alpha 4 | 1.57 | 22.20 |
| *Stc2* | Stanniocalcin 2 | 2.40 | 20.49 |
| *Kif1a* | Kinesin family member 1A | 2.88 | 20.41 |
| *Gm32857* | Predicted gene, 32857 | 3.64 | 20.41 |
| *Col18a1* | Collagen, type XVIII, alpha 1 | 1.05 | 20.29 |
| *Abcc3* | ATP-binding cassette, sub-family C (CFTR/MRP), member 3 | 2.05 | 19.39 |
| *Astn2* | Astrotactin 2 | 2.08 | 19.23 |
| *Slc38a1* | Solute carrier family 38, member 1 | 1.78 | 19.08 |
| *Mthfd2* | Methylenetetrahydrofolate dehydrogenase (NAD+ dependent), methenyltetrahydrofolate cyclohydrolase | 1.89 | 19.08 |
| *Eda2r* | Ectodysplasin A2 receptor | 2.53 | 18.69 |
| *Ugt2b35* | UDP glucuronosyltransferase 2 family, polypeptide B35 | 3.39 | 18.43 |
| *Egf* | Epidermal growth factor | -3.06 | 99.15 |
| *Gpx6* | Glutathione peroxidase 6 | -2.50 | 55.12 |
| *Kcnt1* | Potassium channel, subfamily T, member 1 | -3.94 | 43.24 |
| *Umod* | Uromodulin | -2.09 | 36.46 |
| *Jun* | Jun proto-oncogene | -3.19 | 30.59 |
| *Zfp36* | Zinc finger protein 36 | -2.62 | 27.09 |
| *Btg2* | BTG anti-proliferation factor 2 | -2.90 | 26.64 |
| *Tmem207* | Transmembrane protein 207 | -2.71 | 22.26 |
| *Ckb* | Creatine kinase, brain | -1.84 | 22.20 |
| *Ier2* | Immediate early response 2 | -2.95 | 20.44 |
| *Wfdc15b* | WAP four-disulfide core domain 15B | -1.90 | 20.41 |
| *Nr4a1* | Nuclear receptor subfamily 4, group A, member 1 | -3.63 | 19.45 |
| *Pex5l* | Peroxisomal biogenesis factor 5-like | -2.83 | 19.25 |
| *Fos* | FBJ osteosarcoma oncogene | -3.41 | 17.78 |
| *Ccn1* | Cellular communication network factor 1 | -3.15 | 17.74 |
| *Ppp1r1a* | Protein phosphatase 1, regulatory inhibitor subunit 1A | -1.27 | 17.43 |
| *Snord14e* | Small nucleolar RNA, C/D box 14E | -2.76 | 17.17 |
| *Dusp15* | Dual specificity phosphatase-like 15 | -2.35 | 17.14 |
| *Csrnp1* | Cysteine-serine-rich nuclear protein 1 | -2.46 | 17.07 |
| *Ier3* | Immediate early response 3 | -2.28 | 16.10 |
| *Gm36797* | Predicted gene, 36797 | -2.79 | 15.52 |
| *Nr4a2* | Nuclear receptor subfamily 4, group A, member 2 | -2.96 | 14.57 |
| *Tob1* | Transducer of ErbB-2.1 | -1.65 | 14.36 |
| *Ccn2* | Cellular communication network factor 2 | -1.62 | 14.05 |
| *Gm17971* | Predicted gene, 17971 | -2.05 | 14.02 |

FDR: False discovery rate-adjusted *P* value (Benjamini-Hochberg correction).

**Supplementary Table 12: Primers used for real-time RT-PCR analysis.**

| Gene product | Forward primer (5'-3') | Reverse primer (5'-3') | PCR Product (bp) | Efficiency |
|---|---|---|---|---|
| *18S* | GTA ACC CGT TGA ACC CCA TT | CCA TCC AAT CGG TAG TAG CG | 151 | 0.98 ± 0.02 |
| *36B4* | CTT CAT TGT GGG AGC AGA CA | TTC TCC AGA GCT GGG TTG TT | 150 | 1.02 ± 0.02 |
| *Acox1* | CTG GTG GGT GGT ATG GTG TC | GTG ACT CAC TTG GGC CTG AA | 186 | 1.03 ± 0.03 |
| *Acox2* | AAG CCT CAT CCA ACG TGA CC | AAT GCG TTC AGG ACC GTC TT | 151 | 0.99 ± 0.02 |
| *Acox3* | CAT GTA CGA CTG GTC CCT GG | CCC ATG ACT CAG TTC GGT GA | 160 | 1.02 ± 0.03 |
| *Acta2* | TGT GCT GGA CTC TGG AGA TG | GAA GGA ATA GCC ACG CTC AG | 148 | 1.03 ± 0.02 |
| *Actg1* | TGC CCA TCT ATG AGG GCT AC | CCC GTT CAG TCA GGA TCT TC | 102 | 1.03 ± 0.04 |
| *Adgre1* | CCA GGA GTG GCT TTT GTC TC | GGC TTG GAG AAG TCC TCC TT | 152 | 0.97 ± 0.03 |
| *Atf3* | CCA GGT CTC TGC CTC AGA AG | CCG ATG GCA GAG GTG TTT AT | 151 | 0.98 ± 0.03 |
| *Atf4* | CAT GCC AGA TGA GCT CTT GA | GGC AAC CTG GTC GAC TTT TA | 145 | 0.96 ± 0.03 |
| *Ccnd1* | AGC AGA AGT GCG AAG AGG AG | CAA GGG AAT GGT CTC CTT CA | 149 | 0.98 ± 0.03 |
| *Cd68* | CCA ACA AAA CCA AGG TCC AG | ATT GTA TTC CAC CGC CAT GT | 152 | 1.03 ± 0.03 |
| *Col1a1* | GAT CTC CTG GTG CTG ATG GA | GAC CTT GTT TGC CAG GTT CA | 156 | 0.98 ± 0.03 |
| *Col3a1* | TCC TGG TGG TCC TGG TAC T | TTG CCA GGA GAA CCA CTG TT | 154 | 0.96 ± 0.04 |
| *Cpt1a* | TGG CAG TCG ACT CAC CTT TC | ACA CCA TAG CCG TCA TCA GC | 166 | 0.98 ± 0.02 |
| *Cpt2* | TTG ACG CCA TTC AGT TTC AG | GCA GTG CTG CAG GAT TCA TA | 148 | 1.02 ± 0.03 |
| *Cryab* | ACT TCC CTG AGC CCC TTC TA | CTT GCC GTG GAC CTC AAT CA | 186 | 0.98 ± 0.02 |
| *Dnaja4* | TGA AGG CAT CGG TGG GAA AA | AGT TCT CAC AGC GGT CCT TG | 176 | 1.02 ± 0.03 |
| *Dnajb4* | GAC CCT CCC GTC TCA AAC AA | TGA TTT TGG TGC CTT CTT TCC AC | 195 | 1.01 ± 0.02 |
| *Ddit3* | CCA GGA GGA AGA GGA GGA AG | CCG CTC GTT CTC TTC AGC TA | 148 | 1.02 ± 0.03 |
| *Fn1* | GCA AGC CAG TTT CCA TCA AT | CAT TTT TGG GAG TGG TGG TC | 150 | 0.98 ± 0.02 |
| *Gapdh* | TGC ACC ACC AAC TGC TTA GC | GGA TGC AGG GAT GGG GGA GA | 176 | 1.04 ± 0.03 |
| *Hprt1* | ACA TTG TGG CCC TCT GTG TG | TTA TGT CCC CCG TTG ACT GA | 162 | 0.99 ± 0.01 |
| *Hsp90aa1* | CCC GTG AAA TGC TGC AAC AA | GTA CCG CAA CAG CTC TGA AAG | 200 | 0.98 ± 0.03 |
| *Hsp90ab1* | GAC CTG CCC CTG AAC ATC TC | GGC GTC GGT TAG TGG AAT CT | 196 | 1.04 ± 0.02 |
| *Lcn2* | ATG TCA CCT CCA TCC TGG TC | GTG GCC ACT TGC ACA TTG TA | 148 | 0.97 ± 0.03 |
| *Lgals3* | GCC TAC CCC AGT GCT CCT | TTG CGT TGG GTT TCA CTG TG | 151 | 0.98 ± 0.02 |
| *Mki67* | TGC AAA GGT AGA GGC TCC AT | CAG GTA GGC CAG AGC AAG T | 152 | 0.98 ± 0.03 |
| *Nupr1* | ACC CTT CCC AGC AAC CTC TA | TGG AAC TTG GTC AGC AGC TT | 187 | 0.97 ± 0.02 |
| *Pcna* | TTG GAA TCC CAG AAC AGG AG | ATT GCC AAG CTC TCC ACT TG | 155 | 1.04 ± 0.03 |
| *Ppia* | CGT CTC CTT CGA GCT GTT TG | CCA CCC TGG CAC ATG AAT C | 139 | 1.02 ± 0.02 |
| *Ptprc* | GGA GAC CAG GAA GTC TGT GC | GTT CTG GGC TCC TTC CTC TT | 145 | 0.97 ± 0.03 |
| *Sec61a1* | CGT TGG TGG CCT GTG TTA CT | CAC CAT CTG CTG CTC CTT CA | 190 | 1.02 ± 0.03 |
| *Slc12a1* | ATT GGC CTG AGC GTA GTT GT | AGC AAA GAT CAA GCC TAT TGA CC | 150 | 1.01 ± 0.03 |
| *Tgfb1* | GTG GAA ATC AAC GGG ATC AG | GTT GGT ATC CAG GGC TCT C | 150 | 0.96 ± 0.03 |
| *Tlr4* | GTG GCC CTA CCA AGT CTC AG | GAC CCA TGA AAT TGG CAC TC | 154 | 1.01 ± 0.02 |
| *Umod* | TCA GCC TGA AGA CCT CCC TA | GAA AAG CCT CAG TGG ACA GC | 156 | 0.99 ± 0.02 |

**Supplementary Table 13: RNA-Seq quality and yield.**

| Sample | Genotype (*Umod*) | Age (months) | Yeld (Mbp) | %Q30 | Mean Q |
|---|---|---|---|---|---|
| WT_1.1 | +/+ | 1 | 4'802 | 91.63 | 34.75 |
| WT_1.2 | +/+ | 1 | 6'401 | 90.87 | 34.47 |
| WT_1.3 | +/+ | 1 | 5'373 | 93.09 | 35.16 |
| C171Y_1.1 | C171Y/+ | 1 | 5'937 | 91.76 | 34.82 |
| C171Y_1.2 | C171Y/+ | 1 | 6'435 | 95.13 | 35.73 |
| C171Y_1.3 | C171Y/+ | 1 | 5'301 | 91.87 | 34.83 |
| R186S_1.1 | R186S/+ | 1 | 5'132 | 92.7 | 35.07 |
| R186S_1.2 | R186S/+ | 1 | 4.891 | 92.81 | 35.07 |
| R186S_1.3 | R186S/+ | 1 | 5'212 | 89.28 | 34.22 |
| WT_4.1 | +/+ | 4 | 6'592 | 90.15 | 34.45 |
| WT_4.2 | +/+ | 4 | 7'648 | 91.63 | 34.75 |
| WT_4.3 | +/+ | 4 | 8'959 | 93.77 | 35.42 |
| C171Y_4.1 | C171Y/+ | 4 | 6'932 | 92.41 | 35.01 |
| C171Y_4.2 | C171Y/+ | 4 | 9'851 | 93.28 | 35.36 |
| C171Y_4.3 | C171Y/+ | 4 | 7'213 | 92.23 | 34.97 |
| R186S_4.1 | R186S/+ | 4 | 6'392 | 91.81 | 34.79 |
| R186S_4.2 | R186S/+ | 4 | 6'809 | 91.74 | 34.72 |
| R186S_4.3 | R186S/+ | 4 | 5'311 | 91.77 | 34.98 |

All reads have passed the Illumina chastity filter. %Q30, percentage of bases with quality score ≥30; Mean Q, prediction of the probability of a wrong base call.
